# A computational and multi-brain signature for aberrant social coordination in schizophrenia

**DOI:** 10.1101/2024.02.27.582430

**Authors:** Ya-Jie Wang, Yalan Wen, Leilei Zheng, Ji Chen, Zheng Lin, Yafeng Pan

**Affiliations:** Department of Psychology and Behavioral Sciences, Zhejiang University, Hangzhou, China; Department of Psychiatry, Second Affiliated Hospital, School of Medicine, Zhejiang University, Hangzhou, China; Department of Psychiatry, The Fourth Affiliated Hospital, Zhejiang University School of Medicine, Yiwu, Zhejiang, China

**Keywords:** social coordination, schizophrenia, interpersonal synchronization, hyperscanning, fNIRS, dynamical systems

## Abstract

Social functioning impairment is a core symptom of schizophrenia (SCZ). Yet, the computational and neural mechanisms of social coordination in SCZ under real-time and naturalistic settings are poorly understood. Here, we instructed patients with SCZ to coordinate with a healthy control (HC) in a joint finger-tapping task, during which their brain activity was measured by functional near-infrared spectroscopy simultaneously. The results showed that patients with SCZ exhibited poor rhythm control ability and unstable tapping behaviour, which weakened their interpersonal synchronization when coordinating with HCs. Moreover, the dynamical systems modelling revealed disrupted between-participant coupling when SCZ patients coordinated with HCs. Importantly, increased inter-brain synchronization was identified within SCZ-HC dyads, which positively correlated with behavioural synchronization and successfully predicted dimensions of psychopathology. Our study suggests that SCZ individuals may require stronger neural alignment to compensate for deficiency in their coordination ability. This hyperalignment may be relevant for developing inter-personalized treatment strategies.

## 1 Introduction

Social coordination is one of the most important human social behaviours, playing a pivotal role in human social bonding and mental health^1,2^. Social coordination is defined as the process by which individuals engage in patterned and synchronized behaviours to achieve common goals during social interactions^1,2^, and can be typically characterized by elevated interpersonal synchronization^3^. Despite being a natural and instinctive phenomenon, psychiatric disorders, including schizophrenia (SCZ), often manifest as disruptions in interpersonal coordination^4,5^. For instance, studies have shown that individuals with SCZ exhibited impairments in interpersonal synchronization when coordinating with partners^6,7^.Exploring the manifestations and underlying mechanisms of social coordination deficits in SCZ can yield fresh insights for the development of novel treatment and interventions^8,9^.

A useful approach to addressing the problem of coordination and synchronization involves the use of a phase oscillator model, wherein each member of a group is abstracted as an oscillator^7^. Among the various models of coupled phase oscillators, the Kuramoto model is recognized as a prominent example to capture the synchronization behaviour^10^. Given SCZ is the syndrome of disorganization in behaviour^11,12^, the Kuramoto model can be instrumental in understanding how individuals with SCZ exhibit impairments in organization and coordination. Expanding on the previous research^13^, we performed a formal examination of this concept by adapting a Kuramoto model for group synchronization. This model posits that synchronization emerges as a function of both external and internal coupling strengths^14^. By examining the coupling strengths of SCZ patients coordinating with healthy individuals in diverse scenarios, we are enabled to elucidate the computational processes (i.e., by leveraging dynamical systems) underlying the impaired social functioning observed in SCZ.

Besides the efforts in behavioural sciences and computational modeling, researchers have also examined impairments in social cognition using neuroimaging approaches focused on individual participants^15^. However, it is important to acknowledge that the social brain cannot be fully understood in isolation. The use of hyperscanning techniques (i.e., the simultaneous measurement of two or more brains^16^) provides a promising avenue for studying live interactions between individuals. Recent neuroimaging studies have demonstrated that the neural activities of two individuals become synchronized when they perform a cooperative button-pressing task together^17^. Such inter-brain synchronization (IBS) was closely associated with the level of information exchange between partners^18,19^. Hence, the hyperscanning technique and IBS metric provide a powerful tool for investigating the neural processes associated with atypical social interactions in SCZ^20^.

Currently, the empirical research dissecting the specific role of IBS in SCZ patients, where impaired social functioning represents a core symptom, is sparse. On the one hand, according to the “coordination-by-synchrony” hypothesis^21^, it is reasonable to infer that individuals with SCZ may demonstrate decreased IBS due to deficiency in their coordination ability. Indeed, there is evidence that impairment in social functioning is associated with poor synchrony^8^. In a recent functional near-infrared spectroscopy (fNIRS) hyperscanning study that involved dyads consisting of individuals at clinical high risk of psychosis (CHR) and healthy controls (HC)^22^, reduced behavioural synchrony and IBS in CHR-HC dyads compared to HC-HC dyads was detected. On the other hand, researchers have also observed that IBS, in turn, tends to increase during aberrant social coordination. Consistent with phenomenon, the so-called “compensatory recruitment” hypothesis postulated that social interaction deficits might to some extent require increased neural activities including IBS^23^. Recent research on individuals with social dysfunction, such as Autism Spectrum Disorder, has demonstrated elevated levels of IBS during tasks involving social coordination^24^, suggesting that heightened social interaction demands may contribute to an increase in IBS^25,26^. This pattern may also be applicable to individuals with SCZ, as compensatory adjustments in alignment during interactions may be initiated when dyad members experience deficits in communication^11^. Considering the potential compensatory mechanisms at play, it is plausible that the individuals with SCZ coordinating with others may need a higher level of IBS to successfully carry out a coordination task.

In this study, a total of 114 participants were recruited. The participants were randomly divided into two groups: 35 SCZ-HC dyads and 44 HC-HC dyads (see **Supplementary Table S1** and **S2** for detailed participants information). The goal of the present study was two-fold, the first being to examine the behavioural, computational, and neural mechanisms in SCZ patients coordinating with HCs. To achieve this goal, we used the joint finger-tapping paradigm^27^ (i.e., two persons in a dyad tap an isochronous rhythm together) to assess social coordination, the Kuramoto model for dynamical system modeling, and the fNIRS hyperscanning technique to capture multi-brain neural activities (see **Fig. 1** and **Methods**). Second, we aimed to elucidate how these behavioural and neural features were linked to their behavioural performance and, importantly, the symptomatology of individuals with SCZ. We examined the relationships between IBS and behavioural performance, and sought the IBS patterns that best represented coupling strength derived from dynamical systems modeling. Complementary to this, we used the partial least squares regression (PLSR) to predict dimensions of symptomatology based on IBS.

**Figure 1.**
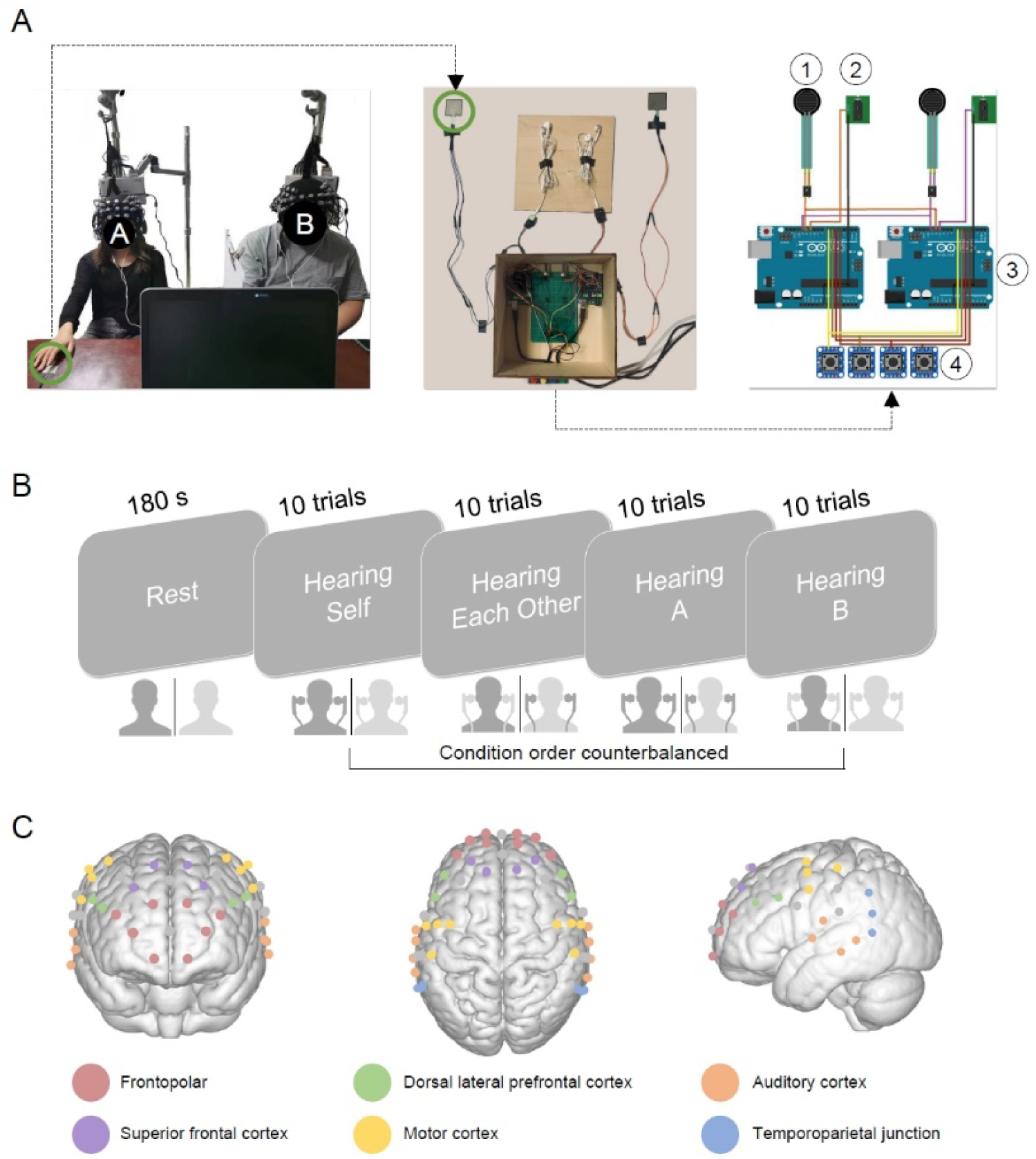
Experimental design. (A) An example of the experimental setup and detailed information about the apparatus. The positions of participants A and B were randomized. In the HC-HC group, roles A and B were assigned randomly to participants, while in the SCZ-HC group, the role B was always a SCZ patient. ① Force-sensitive resistor, ② Headphone jack, ③ Arduino microcontroller, and ④ Condition selection buttons. (B) The experimental procedure and the illustration of the joint finger-tapping paradigm. The headphones’ color indicated the source of auditory feedback (i.e., from self or the partner). The auditory feedback for the dyad can be adjusted to create different coupling conditions: non-coupled, where both members hear only their own beats (Hearing Self); bidirectionally coupled, where each member hears the beats of the other (Hearing Each Other); and unidirectionally coupled, where one member hears the other, and the second member hears only themselves (Hearing A or Hearing B). (C) Localization of channels and regions of interest (ROIs) for fNIRS measurements. Channels of the same color were grouped together to form an ROI.

## 2 Results

### 2.1 Disrupted social coordination in SCZ

As a first-pass analysis, we examined the behavioural performance in SCZ patients when they coordinated with partners. This entailed the evaluation of rhythm deviation (i.e., the absolute deviation from metronome beats set at a standard 500 ms interval), tapping variability (i.e., the stability of coordination), and interpersonal synchronization (i.e., the degree of synchronization within the dyad).

First, the Linear Mixed Model (LMM) results regarding rhythm deviation showed a significant main effect of Role (*F_1,539_* = 24.00, *p <* 0.001, η_p_^2^ = 0.04 (95% confidence interval (CI): 0.02, 1.00)) and an interaction effect for Role × Group (*F*_1,539_ = 15.87, *p* < 0.001, η_p_^2^ = 0.03 (95% CI: 0.01, 1.00)). No other significant main effects or interaction effects were observed. Role A (Median ± median absolute deviation (MAD), 31.50 ± 11.30) showed superior rhythm control ability compared to Role B (34.90 ± 14.30). It is important to note that Role A consistently represents a healthy participant, whereas Role B serves as a healthy participant in HC-HC dyads and as a patient with SCZ in SCZ-HC dyads. Further analysis revealed that all HCs had better beat-keeping ability than individuals with SCZ (44.80 ± 21.90, **Fig. 2A**): HC _Role A_ in the HC-HC dyad (29.40 ± 9.45, *t*_151_ = 6.68, *p* < 0.001, Cohen’s *d* = 0.99 (95% CI: 0.69, 1.28)), HC _Role B_ in the HC-HC dyad (31.60 ± 10.20, *t*_151_ = 6.18, *p* < 0.001, Cohen’s *d* = 0.91 (95% CI: 0.62, 1.21)), and HC _Role A_ in the SCZ-HC dyad (34.50 ± 13.90, *t*_539_ = 5.95, *p* < 0.001, Cohen’s *d* = 0.71 (95% CI: 0.47, 0.95)).

**Figure 2.**
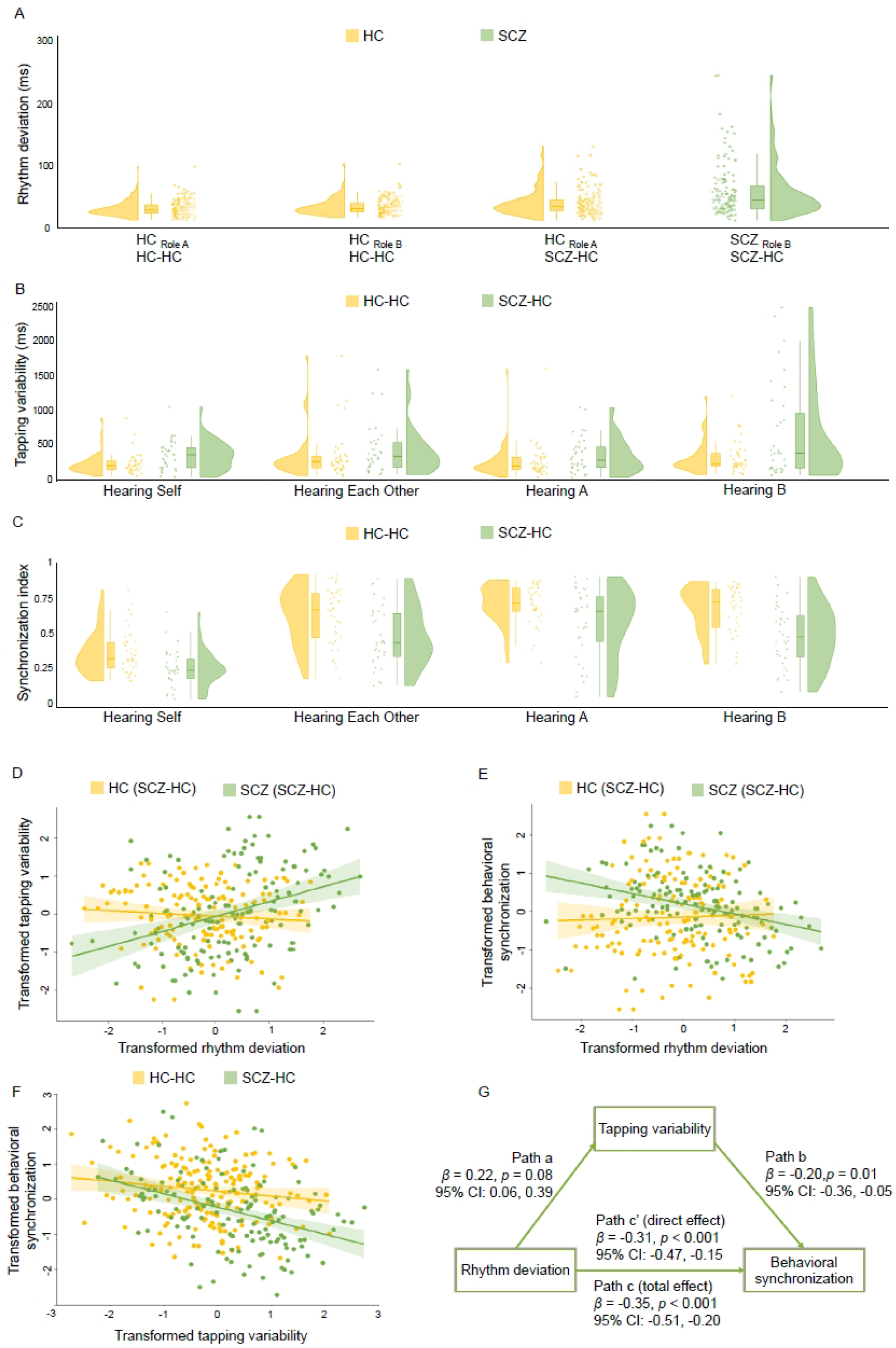
Measures of social coordination. (A) Rhythm deviation across the roles and groups. HC-HC group: role A/B = HC; SCZ-HC group: role A = HC, role B = SCZ. (B) Tapping variability and (C) Synchronization indices across the groups and conditions. (D-F) Pairwise correlation plots for rhythm deviation, tapping variability and behavioural synchronization. All variables in the correlation plots were transformed to conform normal distributions using the Rank-Based Inverse Normal Transformation. (G) The mediation results. The rhythm deviation impacts behavioural synchronization partially through tapping variability.

Second, the LMM analysis on tapping variability revealed significant effects of Group (*F*_1,77_ = 12.28, *p* < 0.001, η_p_^2^ = 0.14 (95% CI: 0.04, 1.00)) and Condition (*F*_3,231_ = 5.85, *p* < 0.001, η_p_^2^ = 0.07 (95% CI: 0.02, 1.00)), with no observed interaction effects (*F*_3,231_ = 2.15, *p* = 0.09). Post-hoc analysis showed that SCZ-HC dyads (371.00 ± 259.00) exhibited more variability in behaviour than HC-HC dyads (233.00 ± 107.00, **Fig. 2B**). Hearing B (286.00 ± 170.00) demonstrated significantly more variability than Hearing Self (248.00 ± 145.00, *t*_231_ = 3.60, *p* = 0.002, Cohen’s *d* = 0.58 (95% CI: 0.26, 0.90)) and Hearing A (247.00 ± 138.00, *t*_231_ = 3.51, *p* = 0.003, Cohen’s *d* = 0.56 (95% CI: 0.24, 0.88)), but showed no difference compared to Hearing Each Other (286.00 ± 174.00, *t*_231_ = 1.68, *p* = 0.33).

Third, in the analysis of the synchronization index, the results of the LMM revealed significant effects of both Group (*F*_1,77_ = 21.94, *p* < 0.001, η_p_^2^ = 0.22 (95% CI: 0.10, 1.00)) and Condition (*F*_3,231_ = 80.39, *p* < 0.001, η_p_^2^ = 0.51 (95% CI: 0.44, 1.00)), with no significant interaction effects (*F*_3,231_ = 1.22, *p* = 0.30). Post-hoc analyses revealed that the SCZ-HC group (0.43 ± 0.26) showed a lower level of synchronization as opposed to the HC-HC group (0.65 ± 0.23). Furthermore, Hearing Self (0.30 ± 0.11) exhibited the weakest synchronization compared to the other three conditions (Hearing Each Other: 0.59 ± 0.24; Hearing A: 0.68 ± 0.19; Hearing B: 0.61 ± 0.23, *t*s > 11.52, *p*s < 0.001, Cohen’s *ds* > 1.70, lower-limited 95% CIs > 1.35, upper-limited 95% CIs > 2.04, **Fig. 2C**). Hearing A had higher synchronization than both Hearing Each Other (*t*_231_ = 3.95, *p* < 0.001, Cohen’s *d* = 0.63 (95% CI: 0.31, 0.95)) and Hearing B conditions (*t*_231_ = 3.03, *p* = 0.01, Cohen’s *d* = 0.49 (95% CI: 0.17, 0.80)), but no significant difference was observed between the synchronization levels of Hearing Each Other and Hearing B (*t*_231_ = 0.92, *p* = 0.79).

We used correlation analysis to explore the relationships among the three variables. In individuals with SCZ, we observed a significant correlation between rhythm deviation and tapping variability (*r* = 0.32, *p*_FDR_ < 0.001), while this correlation was absent in HC individuals (*r* = -0.07, *p*_FDR_ = 0.40, **Fig. 2D**). Conversely, rhythm deviation was negatively correlated with synchronization index only in SCZ individuals (*r* = -0.36, *p*_FDR_ < 0.001), with no such correlation observed in HC individuals (*r* = -0.15, *p*_FDR_ = 0.18, **Fig. 2E**). Similarly, tapping variability and synchronization index displayed a negative correlation in the SCZ-HC group (*r* = -0.40, *p*_FDR_ < 0.001), whereas no such correlation was evident in the HC-HC group (*r*= -0.14, *p*_FDR_ = 0.18, **Fig. 2F**). For additional details on behavioural correlation results in the HC-HC group, please refer to **Supplementary Text** and **Supplementary Fig.S1**.

The mediation model indicated that the rhythm deviation had an impact on behavioural synchronization partially through tapping variability (**Fig. 2G**). Rhythm deviation positively predicted tapping variability (β = 0.22, *p* = 0.08 (95% CI: 0.06, 0.39)) and negatively predicted behavioural synchronization (Direct Effect: β = -0.31, *p* < 0.001 (95% CI: -0.47, -0.15); Total Effect: β = -0.35, *p* < 0.001 (95% CI: -0.51, -0.20)). Additionally, tapping variability negatively predicted the behavioural synchronization (β = -0.20, *p* = 0.01 (95% CI: -0.36, -0.05)). The 95% CI for partially standardized indirect effects was [-0.17, -0.01].

To further uncover the computational basis for social coordination, we used a four-oscillator Kuramoto model to quantify the synchronization patterns^13^. The model abstracts each member of a dyad as a unit comprised of two interconnected oscillators, representing intrinsic processes of perception and action. By applying this model to behavioural data, we could illustrate the relationship between within- (internal, *i*) and between-unit (external, *e*) coupling across different groups and conditions. All the group-level optimal coupling weights were summarized in the **Table 3**. In the Hearing Self condition, both groups showed high internal and low external coupling, suggesting limited social interaction between participants. Conversely, in the Hearing Each Other condition, the coupling patterns reversed, with high external coupling and reduced internal coupling, indicating active social interactions between participants. Within the HC-HC group, asymmetrical external coupling was evident during the Hearing A and Hearing B conditions, indicating unidirectional information feedback. Interestingly, the SCZ-HC group exhibited comparable coupling patterns during the Hearing B condition, but showed attenuated external coupling specifically when hearing A (**Table 3**). The computational modeling results indicate that when hearing HC (i.e., both individuals received auditory feedback from HCs), the SCZ-HC group showed reduced external coupling.

**Table 3.** The optimal internal (*i*) and external (*e*) coupling weights obtained from the four-oscillator Kuramoto modeling.

| Overview of best coupling weights |  | Unit 1 |  | Unit 2 |  |
| --- | --- | --- | --- | --- | --- |
| Coupling term | | $i_1$ | $e_1$ | $i_2$ | $e_2$ |
| HC-HC | Hearing Self | 8.20 | 2.10 | 9.40 | 1.70 |
|  | Hearing Each Other | 8.00 | 9.50 | 10.20 | 10.30 |
|  | Hearing HC<sub>Role A</sub> | 9.10 | 5.70 | 9.40 | 7.90 |
|  | Hearing HC<sub>Role B</sub> | 10.20 | 6.40 | 9.30 | 6.00 |
| SCZ-HC | Hearing Self | 9.50 | 2.30 | 8.80 | 3.10 |
|  | Hearing Each Other | 8.00 | 10.20 | 9.10 | 10.70 |
|  | Hearing HC<sub>Role A</sub> | 9.40 | 2.00 | 9.50 | 2.90 |
|  | Hearing SCZ<sub>Role B</sub> | 9.20 | 5.80 | 9.40 | 4.80 |

### 2.2 Alterations in the multi-brain network during social coordination in SCZ

Having confirmed the social coordination deficits in SCZ, we next investigated the inter-brain activities underpinning these deficits. Before presenting all inter-brain results (including IBS and inter-brain Granger Causality, GC), however, an investigation of intra-brain activation and synchronization is needed as a sanity check.

First, we found that brain activation did not show any significant main effects for Group, Condition, or any interaction effects on any of the ROIs (all *p*_FDR_ > 0.05).

Second, we observed main effects of Group on intra-brain synchronization. Specifically, all nine combinations of ROIs showed significantly lower intra-brain synchronization in the SCZ-HC group compared to the HC-HC group (see **Fig. 3A, Supplementary Text, Supplementary Fig.S3,** and **Supplementary Table S6**). Meanwhile, we observed interaction effects of Group and Role. All four combinations of ROIs consistently showed that SCZ participants in the SCZ-HC dyads showed lower intra-brain synchronization than healthy participants (see **Fig. 3C**, **Supplementary Text, Supplementary Fig.S3,** and **Supplementary Table S6**). No significant main effects of Condition or other interaction effects were found for any ROI. Furthermore, we found significant main effects of Group on intra-brain GC, where 9 out of 11 combinations of ROIs exhibited reduced GC in the SCZ-HC group (see **Figs. 3B** and **3D, Supplementary Text, Supplementary Fig.S4,** and **Supplementary Table S7**).

**Figure 3.**
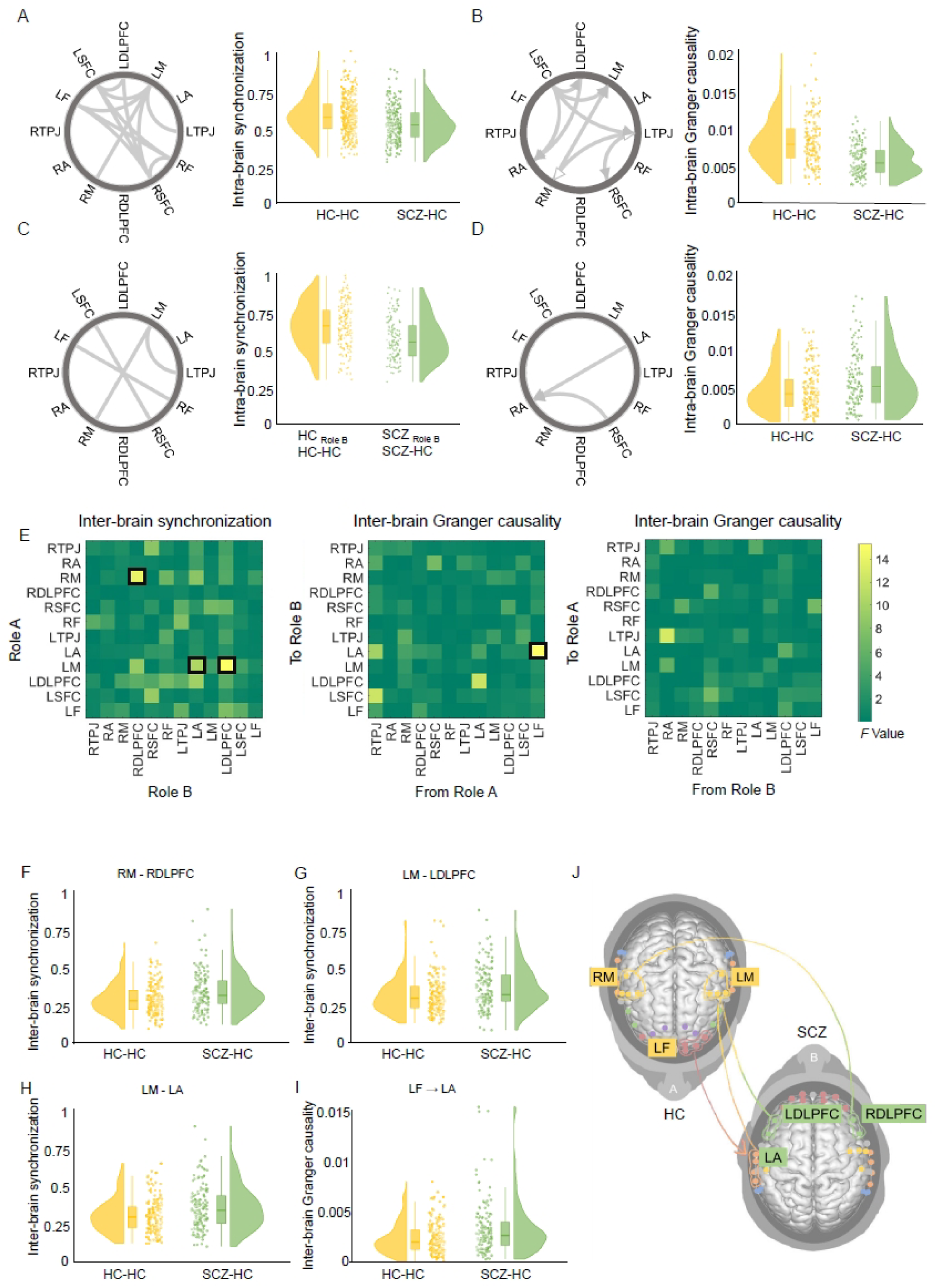
Multi-brain networks. **Intra-brain network.** (A) Regions of interest (ROI) combinations demonstrating significant Group effects on intra-brain synchronizations. (B) ROI combinations indicating significant Group effects on intra-brain Granger Causality (GC), with SCZ-HC dyads showing lower GCs than HC-HC dyads. The black arrow denotes unidirectionality, while the white arrow indicates bidirectionality. (C) ROI combinations showing significant Group × Role interaction effects on intra-brain synchronizations. (D) ROI combinations indicating significant Group effect on intra-brain GCs, with SCZ-HC dyads showing higher GCs than HC-HC dyads. **Inter-brain network.** (E) The *F*-value map of inter-brain synchronization (IBS) (left panel), GC_A to B_ (middle panel), and GC_B to A_ (right panel) among different inter-brain ROI combinations. (F-I) Data distribution for four specific inter-brain ROI combinations (RM – RDLPFC, LM – LDLPFC, LM – LA, and LF → LA), which showed significant group differences. (J) The virtual brains visualize the interacting partners and illustrate the increased IBS and GC.

Third, the results of the LMM on IBS identified three combinations of ROIs that exhibited significant group differences between the HC-HC group and the SCZ-HC group (**Fig. 3E left panel, F-H, J**): (1) RM_HC_ – RDLPFC_SCZ_, *F*_1, 291.61_ = 13.85, *p*_FDR_ = 0.01, η_p_^2^ = 0.05 (95% CI: 0.01, 1.00); (2) LM_HC_ – LA_SCZ_, *F*_1, 294.66_ = 10.28, *p*_FDR_ = 0.046, η_p_^2^ = 0.03 (95% CI: 0.01, 1.00); and (3) LM_HC_ – LDLPFC_SCZ_, *F*_1, 290.53_ = 15.03, *p*_FDR_ = 0.008, η_p_^2^ = 0.05 (95% CI: 0.02, 1.00). These findings indicate that the SCZ-HC group consistently exhibited higher IBS compared to the HC-HC group: 0.31 ± 0.12, 0.33 ± 0.13, 0.31 ± 0.13 in the SCZ-HC group and 0.27 ± 0.09, 0.29 ± 0.10, 0.28 ± 0.10 in the HC-HC group, respectively. Additionally, we consistently observed the strongest IBS at 11 combinations of ROIs under the Hearing B condition (see **Supplementary Text, Supplementary Fig.S2,** and **Supplementary Table S5** for more details).

Fourth, we observed that the inter-brain GC value, from LF_HC_ to LA_SCZ_, was significantly higher in the SCZ-HC group compared to the HC-HC group, *F*_1, 295.90_ = 15.33, *p*_FDR_ = 0.002, η_p_^2^ = 0.05 (95% CI: 0.02, 1.00), 0.0025 ± 0.0017 in the SCZ-HC group and 0.0019 ± 0.0012 in the HC-HC group (**Figs. 3E middle** and **right panels, I, J**).

### 2.3 Associations of the multi-brain network with behavioural performance

We examined associations of multi-brain network that showed significant group effects with behavioural performance. We found that only IBS at LM_HC_ – LDLPFC_SCZ_ showed a significant correlation with synchronization index in the SCZ-HC group, *r* = 0.32, *p*_FDR_ = 0.0006. Such a relationship was absent in the HC-HC group, *r* = -0.09, *p* = 0.22 (**Fig. 4A**). We further explored the relationship between inter-brain and intra-brain synchronization. In the SCZ-HC group, we found a positive correlation between intra-brain synchronization at RSFC_SCZ_ – LSFC_SCZ_, and IBS at RM_HC_ – RDLPFC_SCZ_: *r* = 0.32, *p*_FDR_ = 0.005 (**Fig. 4B**).

**Figure 4.**
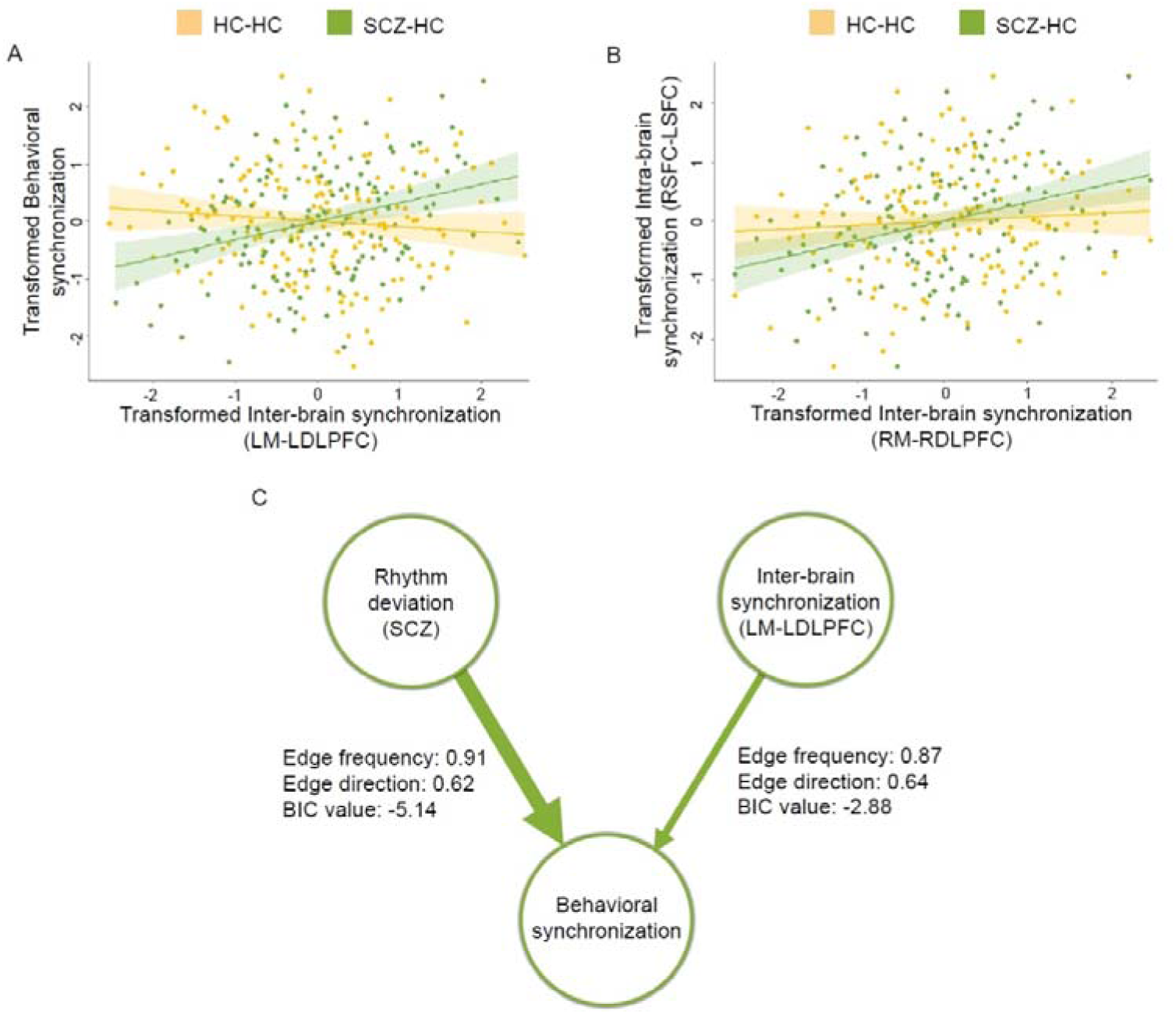
Brain-behaviour relationships. (A) The correlation between inter-brain synchronization (IBS) at LM – LDLPFC and behavioural synchronization. (B) The correlation between IBS at RM – RDLPFC and intra-brain synchronization at RSFC – LSFC. The data were transformed to follow a normal distribution using Rank-Based Inverse Normal Transformation. (C) The Bayesian network results. In the SCZ-HC group, rhythm deviation negatively predicted behavioural synchronization, whereas IBS at LM – LDLPFC positively predicted behavioural synchronization. Variables that failed to engage in this network were omitted in this graph.

A Bayesian network allowed us to explore the directional dependencies among three behavioural indices (rhythm deviation, tapping variability, and behavioural synchronization) and IBS simultaneously, without making explicit assumptions about their relationships. Bayesian networks were used to generate a directed acyclic graph structure, using the hill-climbing algorithm^28^. Using Bayesian network analysis, we found that rhythm deviation could negatively predict behavioural synchronization, and IBS at LM_HC_ – LDLPFC_SCZ_ could positively predict behavioural synchronization in the SCZ-HC group (**Figs. 4C**).

### 2.4 Associations of IBS with external coupling weights derived from dynamical systems modeling

In the dynamical system modeling analysis, we discovered that the most deviated external coupling occurred under the Hearing A condition. We next sought to explore how IBS represented these abnormal external coupling weights using the Mahalanobis distance approach. The results revealed that IBS at RDLPFC-RTPJ best represented both *e*_1_ and *e*_2_ in the HC-HC group, with minimal Mahalanobis distance (i.e., maximal similarity) between IBS and computational parameters. In contrast, in the SCZ-HC group, IBS at LA-RTPJ represented *e*_1_, while IBS at RDLPFC-LSFC encoded *e*_2_ (**Fig. 5**). The full Mahalanobis distance results for the four conditions in both groups can be found in **Supplementary Table S8.**

**Figure 5.**
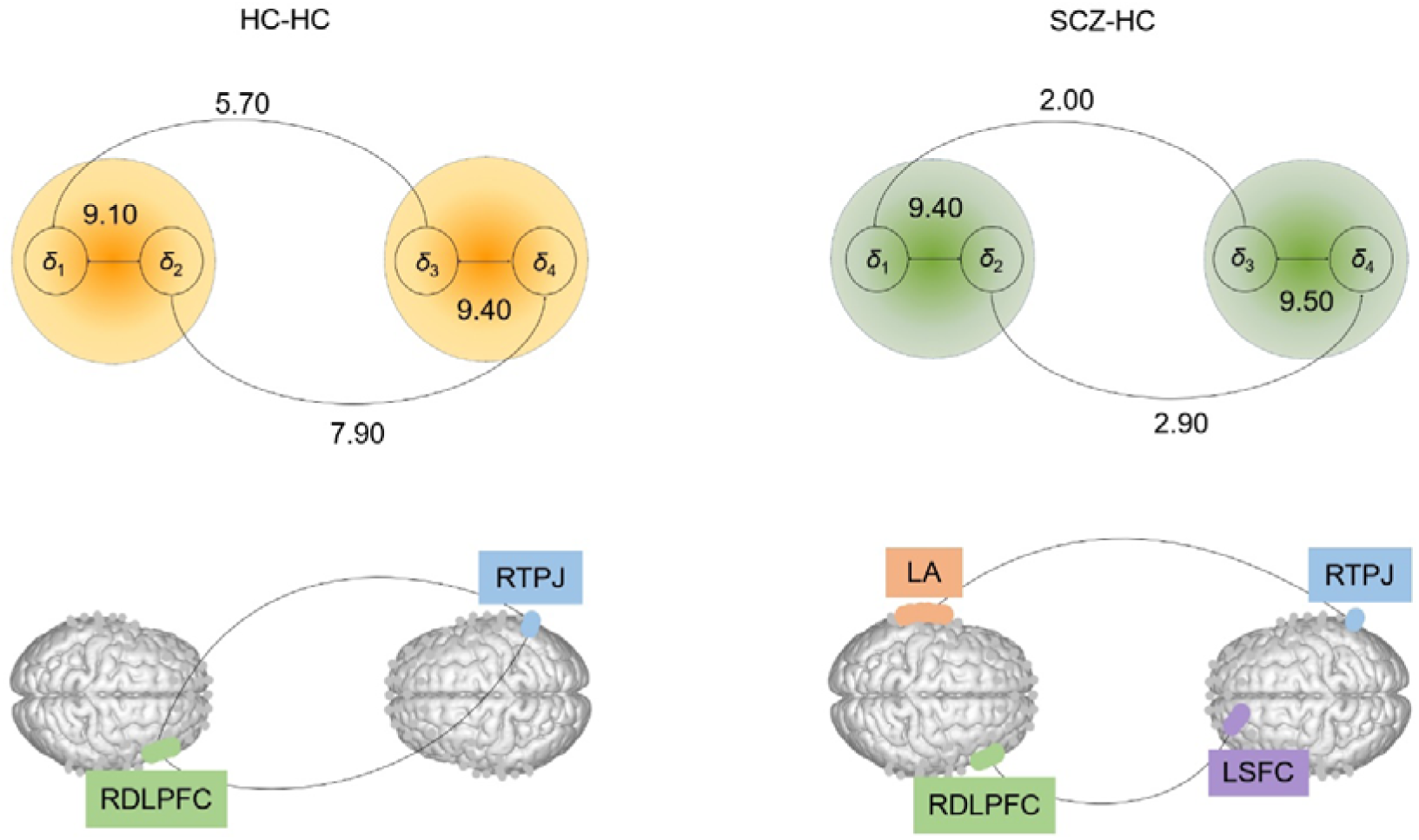
Associations between external coupling weights derived from dynamical systems modeling and inter-brain synchronization.

### 2.5 Predicting dimensions of psychopathology from IBS

Finally, we constructed PLSR models to predict scores of the four symptom dimensions based on IBS. The results showed that IBS was sufficient to predict Total PANSS Score: *r* = 0.58, *p* < 0.001, *R*^2^ = 0.34; Negative symptom: *r* = 0.58, *p* < 0.001, *R*^2^ = 0.33; Positive symptom: *r* = 0.68, *p* < 0.001, *R*^2^ = 0.46; Affective symptom: *r* = 0.54, *p* = 0.001, *R*^2^ = 0.29; and Cognitive symptom: *r* = 0.74, *p* < 0.001, *R*^2^ = 0.54. **Figure 6** illustrates the top three ROI combinations that made the most significant contributions to the latent variable.

**Figure 6.**
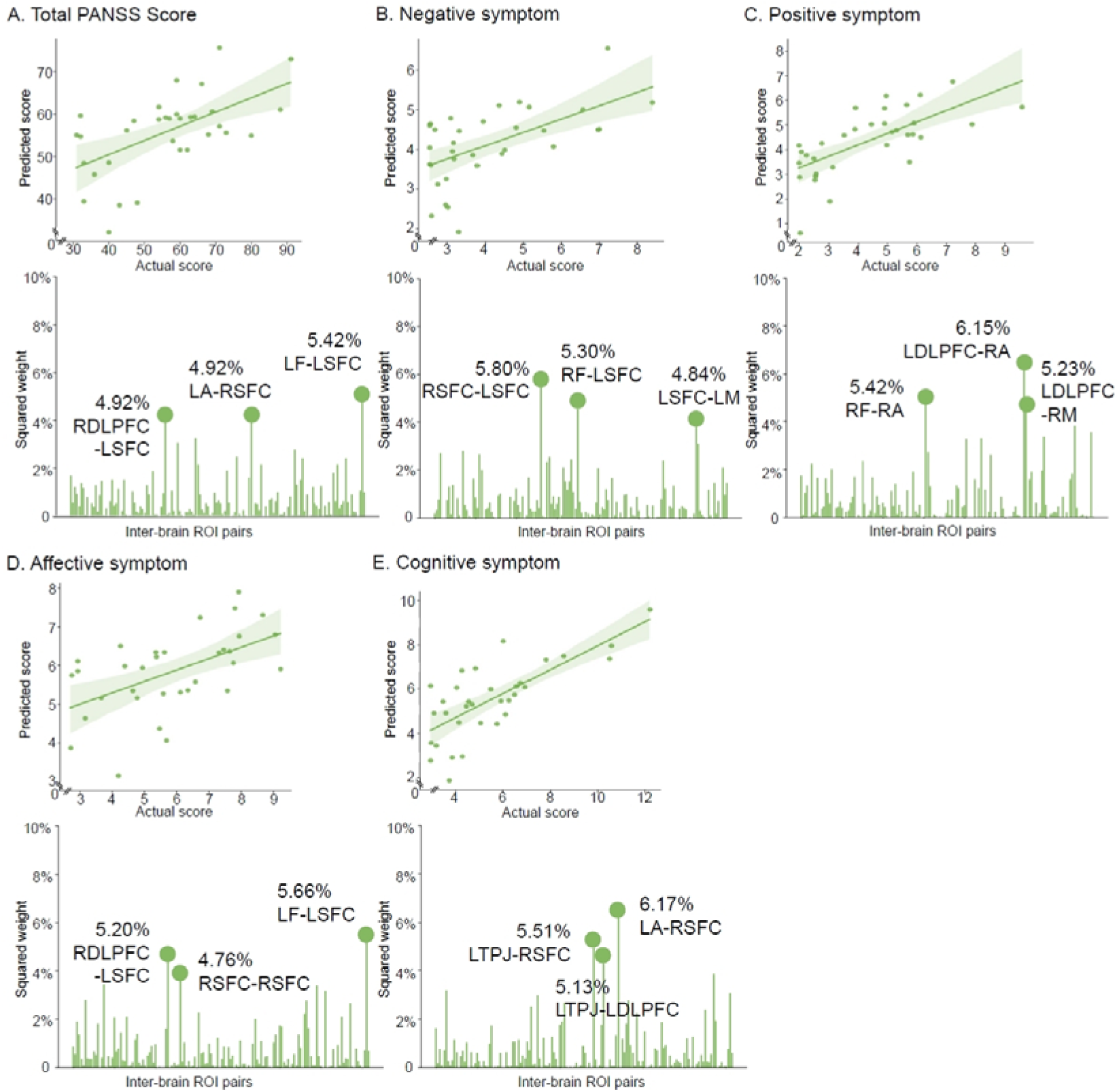
The scatter plots between actual value and predicted values and the squared weights of inter-brain ROI combinations in (A) Total PANSS Score, and scores of (B) Negative symptom, (C) Positive symptom, (D) Affective symptom and (E) Cognitive symptom.

## 3 Discussion

Impairment in interpersonal synchronization is a pervasive characteristic observed in social interactions across various mental disorders, including SCZ^8,29^. This study represents an important undertaking in understanding the disrupted behavioural synchronization and aberrant multi-brain networks in patients with SCZ, within the context of real-time social coordination. The findings revealed that individuals diagnosed with SCZ displayed diminished rhythm control ability and unstable tapping behaviour, leading to reduced interpersonal synchronization while coordinating with HCs. Additionally, we observed decreased external coupling in dyadic dynamical systems when the healthy partner was unable to perceive auditory feedbacks from SCZ. We discovered a decrease in both functional and effective intra-brain connectivity in individuals with SCZ. Importantly, we observed heightened IBS in SCZ-HC dyads, which positively correlated with behavioural synchronization and served as a predictive factor for both behavioural synchronization and symptom severity (measured by PANSS scores).

*First*, alike previous study, we found SCZ patient showed less successful coordinated interactions^8^. Forwardly, our study revealed that less controllable behaviour would hinder interpersonal synchronization partially through the unstable behaviour. Our study support the notion that distributed motor production would lead to imprecise perception^30^, making it difficult for a healthy partner to tune in. Building upon the Kuramoto model, we observed diminished external coupling when in the SCZ-HC dyad when both participants sorely received auditory feedbacks from the HC. In contrast, our observations revealed that the coupling weights of SCZ-HC and HC-HC dyads were comparable in the Hearing B and Hearing Each Other conditions. It is important to note that in these conditions, HC could receive the auditory feedback from individuals with SCZ. However, when HC could only hear their own tapping sound, specifically in the Hearing Self and Hearing A conditions, they were incapable to adjust and coordinate their behaviour as the auditory feedback from individuals with SCZ was absent. Together, these results together suggest that the “normalcy” of these conditions primarily relies on the presence of HC. The lack of accommodation from HC might result in lower external coupling weights and difficulties in achieving overall behavioural coordination. The low external coupling exhibited by the SCZ-HC group may reflect SCZ patients’ deficit in self-other integration in real-time social interactions. This finding echoes earlier research on individual brain functioning, which suggests that SCZ patients face severe difficulties in integrating self and other, leading to problems in social interactions^31–33^. Overall, we found an imbalance pattern on internal and external coupling weights in the SCZ-HC dyads, suggesting a socio-cognitive bias where SCZ preferentially allocate resources to internal rather than external coupling during interactions. This observation supports the notion that the internal model plays a pivotal role in perceptual inference within the SCZ population^34^. Within the Bayesian framework of predictive processing, the problem of disorganization in SCZ can be attributed to a failure to prioritize the shared model and an increased reliance on lower-order internal models^11,30^.

*Second*, we observed that IBS and inter-brain GC were higher in the SCZ-HC dyads compared to the HC-HC dyads, while most intra-brain connectivity was lower in the SCZ-HC dyads. These results on intra-brain connectivity are consistent with previous studies that have suggested a significant role of brain disconnection in the pathophysiology of SCZ^35^. It is important to emphasize that the observed reduction in intra-brain synchronization does not conflict with the previously mentioned concept of “internal coupling strength”. The tendency to allocate resources more to internal coupling than external coupling can be characterized as a disturbance in predictive mechanisms during interpersonal interactions^36^. This phenomenon may be linked to the dysregulation of neural synchronization^34^. However, previous findings on IBS in social deficits are inconsistent, with some reporting an increase^24^, while others report a decrease^37^. Our results provided support for the former, in the SCZ population. Four ROI combinations (i.e., RM–RDLPFC, LM–LDLPFC, LM–LA, and LF→LA) were identified, with consistently higher levels of IBS or inter-brain GC in the SCZ-HC dyad compared to the HC-HC dyad. Considering the functional meaning of the involved brain regions, it is reasonable to observe the involvement of motor and auditory regions in these findings, given that participants received auditory feedback and engaged in button pressing during the task. Regarding DLPFC, this region has been implicated in theory of mind^38,39^. Specific to SCZ, studies have found a negative correlation between DLPFC activity and the severity of negative symptoms, as well as a positive correlation with their level of social functioning^40^. Overall, on the IBS level, HC participants predominantly recruited the motor region, indicating their focus on finger tapping. In contrast, individuals with SCZ exhibited engagement of the auditory region and DLPFC, potentially indicating their effort to receive auditory feedback from the healthy counterparts. This suggests that interpersonal coordination tasks may pose a higher social demand for individuals with SCZ. Our GCA revealed an increased flow of information from the LF_HC_ to the LA_SCZ_. The frontopolar area has been found to have a crucial role in social functioning, as reduced activation in this region is associated with lower social functioning scores^41^. Furthermore, increased IBS at the frontopolar area has been linked to improved communication and coordination performance^42–44^. The frontopolar cortex plays a crucial role in prediction and is actively involved in switching behaviours^45^. It tracks and evaluates the accumulated evidence favoring a switch to an alternative course of action^46^. The recruitment of the frontopolar area in HC suggests that they may have implicitly accommodate SCZ patients in achieving finger tapping synchrony. This information flow ultimately reached the LA of SCZ, indicating their acceptance and processing of auditory signals. Taken together, our findings on the multi-brain network offer a functional understanding of the synchronization phenomenon, indicating that HC might implicitly accommodate with SCZ patients to achieve a state of synchronization.

*Third*, we observed a positive correlation between the intra-brain synchronization (RSFC_SCZ_ – LSFC_SCZ_) and the IBS (RM_HC_ – RDLPFC_SCZ_). Previous study reported that the interpersonal synchronization of SFC was related to social cooperation^17^, suggesting a role in modeling and predicting the behaviour of others. This suggests that the capacity to anticipate the actions of others is closely intertwined with the execution and regulation of motor movements, with the RDLPFC playing a pivotal role in task-related behaviour and the integration of information into dynamic behavioural sequences^47,48^. Moreover, this correlation between intra-brain and inter-brain activity underscores the significance of considering the two brains as an interconnected system. It suggests that individuals with SCZ were not operating in isolation during social coordination tasks; instead, their brains were dynamically adapting to facilitate communication with others. These findings strongly support the concept of a “hyperbrain network”^49,50^, and provide compelling evidence that studying social interaction as a system, rather than in isolation, is essential^51^. An intriguing finding of our study is that the correlation between intra- and inter-brain synchronization was observed exclusively in the SCZ-HC group, rather than the HC-HC group. One plausible explanation for this discrepancy is that the coordination task might be relatively straightforward for HC individuals, thereby not necessitating additional cognitive resources. From this perspective, the observed synchronization could potentially serve as a temporary adaptive mechanism that enables individuals with SCZ or other impairments in social functioning to actively participate in social coordination.

Importantly, our Bayesian network analysis unveiled two robust indicators of behavioural synchronization: rhythm deviation and IBS (LM_HC_-LDLPFC_SCZ_). It is notable that rhythm deviation dampened behavioural synchronization, whereas IBS facilitated it. This underscores the potential significance of IBS as a target for enhancing the recovery of social coordination in individuals with SCZ. Moving forward, future studies could explore IBS as a promising intervention entry point, thereby contributing to a deeper understanding of and potential improvements in SCZ social coordination.

*Forth,* when associating IBS with computational modeling parameters, our model-based findings reveal that SCZ-HC dyads necessitate the engagement of more brain regions compared to HC-HC dyads when both participants can only receive auditory feedback from HC. Notably, the involvement of auditory areas in SCZ-HC dyads supports the hypothesis that SCZ may manifest rigid synchronization patterns from higher to lower-level cortical areas during perception and prediction processes, resulting in imbalanced resource allocation^34^. This observation might reflect the different hierarchies or levels of coordination strategies employed by the SCZ-HC and HC-HC groups.

*Finally*, our analyses revealed a significant correlation between the IBS at LM_HC_–LDLPFC_SCZ_ and behavioural synchronization. Interestingly, despite the lower behavioural synchronization observed in the SCZ-HC dyad, higher levels of IBS were associated with greater behavioural synchronization. This suggests that IBS may act as a compensatory mechanism, aiding individuals with SCZ who have impaired social coordination in achieving better performance. We did not observe an association between intra-brain synchronization and behavioural performance – this highlights the significance of adopting a two-person approach in studying social coordination in SCZ. Our Bayesian network analysis revealed that IBS patterns can serve as a predictor for symptom severity. Among all inter-brain ROI combinations, we extracted the relative importance of their contributions to latent variables that can predict symptoms. Notably, our findings indicate that the previously discussed ROIs, including the SFC, DLPFC, frontopolar, and auditory area, assume relatively more crucial roles in symptom prediction. These findings support the notion that deficits in social coordination could serve as an early indicator of SCZ^22,52–55^. Identifying such markers could facilitate earlier diagnosis and open up new avenues for treatment interventions, such as interpersonal psychotherapy^56^.

In summary, our findings indicate that individuals with SCZ exhibit weakened rhythm control ability, unstable tapping behaviour and poor behavioural synchronization during interpersonal coordination. The use of dynamical systems modeling, specifically the Kuramoto model, further reveals that the support from HC may be crucial for SCZ patients to attain a coherent state. The results from inter-brain network analysis imply the involvement of both partners in coordination: HC may implicitly accommodate with SCZ patients, while the latter may need additional compensatory mechanisms for successful engagement in social coordination. Notably, our results demonstrate that IBS served as compensatory mechanisms for improve behavioural synchronization in SCZ-HC dyads. The predictive power of our inter-brain network for SCZ symptom scores holds promise for potential clinical applications. Overall, our study provides valuable insights into the behavioural, computational, and neural mechanisms underlying social function deficits in SCZ, highlighting the importance of using real-time social tasks and hyperscanning techniques to investigate (aberrant) social interactions.

## 4 Methods

### 4.1 Participants

Participant recruitment and data collection took place between July and December 2022. This study involved 44 dyads in the healthy control group (HC-HC group), consisting of 15 female-female dyads, 15 female-male dyads, and 14 male-male dyads. Each dyad comprised two healthy individuals. Furthermore, there were 35 dyads in the schizophrenia group (SCZ-HC group), including 13 female-female dyads, 13 female-male dyads, and 9 male-male dyads. Each dyad in the SCZ-HC group consisted of one healthy individual paired with one SCZ patient. We calculated the average and absolute differences in age within each dyad and reported the median and MAD to account for non-normality. The median age-average for the HC-HC group was 22.50 years (MAD = 3.71), while the median age for the SCZ-HC group was 24.50 years (MAD = 7.41). In both the HC-HC and SCZ-HC groups, the median age-difference in each dyad was 1.00 year (MAD = 1.48). The two groups were matched in terms of gender and age (*W* = 723, *p* = 0.63 for pair gender, *W*= 888, *p* = 0.24 for age-average, and *W* = 881, *p* = 0.25 for age-difference; see **Supplementary Table S1** and **S2** for detailed demographic information).

The healthy individuals were recruited through various channels, including campus forums, online platforms, and word of mouth. Prospective participants were required to complete the Symptom Checklist-90 (SCL-90) and Edinburgh Handedness Questionnaire as part of their registration process^57,58^. Applicants who scored below 160 on the total SCL-90 scale and below 2.5 on any of the factors^59^, and had a right-handedness score greater than 0.75^60^, were shortlisted. Shortlisted candidates then completed the Fast Symptom Questionnaire for Mental Illness Screening (FSQ-MIS), a newly developed questionnaire specifically designed to screen and assess mental symptoms during the COVID-19 pandemic, with a total score of less than 3 indicating no mental illness^61^ (**Supplementary Table S4**). Additionally, they underwent a psychological assessment conducted by a clinical psychologist. Only participants who did not meet the Diagnostic and Statistical Manual of Mental Disorders, Fifth Edition (DSM-5) diagnostic criteria for any mental disorder were included in the study.

Comprehensive and professional assessments of the SCZ patients were conducted by experienced psychiatrists from the Department of Psychiatry at the Second Affiliated Hospital of Zhejiang University referring to DSM-5. These assessments led to the diagnosis of various schizophrenia spectrum disorders, such as schizophrenia (F20.9), schizoaffective disorder with bipolar (F25.0) and depressive (F25.1) subtypes.

All participants included in this study had no record of significant head trauma, intellectual disability, alcohol abuse, or long-term medication use, except for antipsychotic medications prescribed to patients in the SCZ-HC group. Moreover, participants were free of brain tumors, diabetes, hypertension, cardiovascular diseases, and other severe physical ailments. They had no history of conditions such as transient ischemic attack, hysteria, migraines, syncope, or any other disorders that could potentially impact brain function. Furthermore, participants were not diagnosed with any neurological disorders.

The study received ethical approval from the Research Ethics Committee of the Second Affiliated Hospital of Zhejiang University School of Medicine and adhered to the principles outlined in the Declaration of Helsinki. All participants were informed about the research objectives and provided written consent. Participants were paid for their participation.

### 4.2 Experimental design and procedures

Dyads engaged in a joint finger-tapping task^27^. This task has been widely used to evaluate social coordination between individuals^62,63^. The task was carried out using Tap Arduino (**Fig. 1A**), an Arduino microcontroller designed to provide immediate auditory feedback with minimal delay^64^. A full description of the apparatus can be found in **Supplementary Text**. Two participants in a dyad were positioned on one side of a table, separated by a partition to prevent visual contact and feedback. The experimenter, situated across the participants, controlled the apparatus to regulate the auditory feedback they received (depending on the experimental condition). Each participant used their right index fingers to apply pressure on sensor pads.

After a 180-s rest, dyads underwent four distinct experimental conditions, each having 10 trials (order randomized, **Fig. 1B**). Each trial began with 8 beats at a tempo of 120 beats per minute (bpm). Following this, the stimulus ceased, and the participant received auditory feedback from one of the two sources: their own tapping, their partner’s tapping. The participants were assigned to four different scenarios: (1) Hearing Self — both participants exclusively heard beats generated by themselves; (2) Hearing Each Other — both participants heard beats generated by each other; (3) Hearing A — both participants only heard beats generated by Participant A; and (4) Hearing B — both participants only heard beats generated by Participant B. A stop signal was automatically triggered after a total of 32 button presses, signifying the end of the trial. In the HC-HC group, participants were randomly assigned the roles A or B; in the SCZ-HC group, participant B consistently assumed the role of B (and thus role A was always a healthy individual). The spatial positions of participant A and B were also randomized.

### 4.3 Subjective measurements

*Positive and Negative Syndrome Scale.* The data of the Positive and Negative Syndrome Scale (PANSS) was collected through evaluation by professional psychiatrist within one week of the experimental date. The PANSS is widely recognized as one of the most rigorously validated instruments for assessing positive, negative, and overall psychopathology associated with schizophrenia^65^. This standardized clinical interview examines the presence and severity of positive and negative symptoms, as well as general psychopathology, in individuals diagnosed with schizophrenia over the course of the preceding week. The PANSS consists of 30 items, and the severity of each item is rated on a 7-point scale, with endpoints ranging from 1 (absent) to 7 (extreme). The PANSS 30 items were assigned into four dimensions representing positive, negative, affective and cognitive symptoms of schizophrenia as in our previous studies^66,67^. The detailed information about the contained items for each dimension can be found on **Supplementary Table S4**. Importantly, this 4-dimensional representation of schizophrenia psychopathology is identified to be stable and generalizable across populations, regions and clinical settings. Moreover, the international consistency of this 4-dimensional structure is higher than the PANSS 3 subscales and previously proposed factor structures^66^. Calculation of scores of the four dimensions was performed through an online dimensions and clustering tool that provides assessments of symptomatology for SCZ patients: http://webtools.inm7.de/sczDCTS/. Higher scores denote the more severe symptoms a patient express along particular dimensions.

We also collected *Interpersonal Reactivity Index*, *Schizophrenia Quality of Life Scale*, and *Liebowitz Social Anxiety Scale*. The descriptions for these measurements can be found in **Supplementary Text**. All the results of subjective measurements can be found in **Supplementary Table S1**.

### 4.4 fNIRS data acquisition

Two NirSmart-6000A equipment (Danyang Huichuang Medical Equipment Co., Ltd., China) were used to capture continuous wave fNIRS data throughout the experiment. Each device comprised 21 sources and 16 detectors, resulting in a total of 48 channels (**Fig. 1C**). The source-detector separation distance was approximately 3 cm. To ensure consistency across variations in head size, optodes were placed over standard 10-20 system locations^68^. The virtual registration method was employed to establish the correspondence between the NIRS channels and the measured points on the cerebral cortex^69,70^. The system recorded the absorption of near-infrared light at two wavelengths, specifically 730 and 850 nm, at a sampling rate of 11 Hz. The acquired light absorption data were converted into concentrations of oxy-hemoglobin (HbO) and deoxy-hemoglobin (HbR) using a modified Beer–Lambert law^71^, with a differential pathlength factor of 6^72^. Due to the superior signal-to-noise ratio of HbO compared to HbR^73^, we focused on HbO in the subsequent analyses. At the beginning of the data acquisition process, participants received instructions to maintain a relaxed and awake state while keeping their eyes closed for a duration of 3 minutes. This period was designated for the collection of resting-state fNIRS data.

### 4.5 Data analyses

#### 4.5.1 Behavioural indices

*Rhythm deviation.* This ability was operationalized as the absolute difference between the inter-tap interval (ITI) observed in the real data and the ITI of the computer metronome. The larger absolute difference indicated greater deviation from given beats. As this index exhibited a non-normal distribution, we reported the median value as the representative measure.

*Tapping variability.* This index refers to the level of inconsistency or variation in the timing of finger taps between two partners. In the previous study^62^, tapping variability was quantified using the Standard Deviation (SD) of the asynchronies. Asynchrony represents the time difference between the tap timings of participant A (serving as the reference) and participant B, measured in milliseconds. Due to non-normality, we calculated asynchrony based on each trial, and then the MAD of the absolute value of asynchrony to obtain a measure of finger tapping variability. A higher MAD value suggests a wider adjustment range and less stable behaviour during the tapping task.

*Synchronization index.* To capture the social coordination within dyads, we employed interpersonal synchronization of each participant’s ITI, as a proxy. Synchronization index (SI) was calculated for each participant using the variance of relative phase, following the formula below^74^:

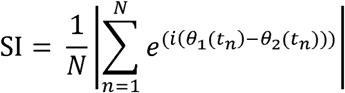

*N* represents the total number of time points, and *n* is each of these time points. *e* denotes Euler’s number (a mathematical constant), *i* stands for the imaginary unit, and θ*_1_*(*t_n_*) signifies the phase at time *n* for the first participant. Consequently, (θ*_1_*(*t_n_*) - θ*_2_*(*t_n_*)) refers to the phase difference between the two participants at a specific time point. The resulting index is a dimensionless measure that ranges from 0 to 1, indicating the degree of synchronization between two interacting agents. If the value exceeds 0.73, it is typically regarded as an indication of synchronous behaviour^75^.

#### 4.5.2 Computational modeling

Behavioural data, specifically ITI, were further parsed using computational modeling. We examined the ability of a four-oscillator Kuramoto model to determine its ability to achieve and sustain synchronization within the limited timeframe observed in joint finger tapping experiments. We specifically selected this type of continuously coupled model due to (1) its extensively documented behaviour, (2) previous successful applications in interpersonal synchronization tasks, and (3) its applicability to a wide range of synchronization phenomena observed in various natural systems^10,13,14,76^. Our aim was to establish a dynamic representation based on social coordination performance, which captured by coupling strength, in both SCZ-HC and HC-HC groups. The Kuramoto model can be described as follows^13,76,77^:

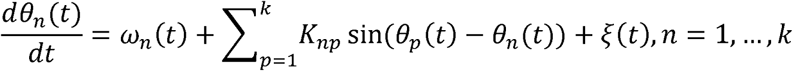

The symbol ω represents the natural frequency of the oscillator. The coupling strengths or the coupling matrix between oscillator *n* and oscillator *p* are denoted by *K_np_*. The variable θ*_n_(t)* represents the phase of oscillator *n* at time *t*. Additionally, the symbol ξ represents a Gaussian noise component that introduces stochasticity into the system and affects its dynamics.

The numerical integration of the Kuramoto model of coupled oscillators was performed using custom codes in MATLAB 2022a, following the methodology described in a previous study^13^. Our implementation utilized the complete four-oscillator model, with the coupling matrix *K_np_* defined according to the following specifications:

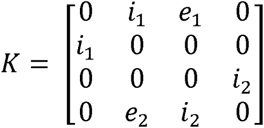

We assigned symbols *i* and *e* to represent the weights of the internal and external coupling, respectively. For the remaining parameters, we set the value of ω to 2 Hz, corresponding to the tapping rate required by the task. The noise component ξ was modeled as a Gaussian distribution with a mean of zero. The standard deviation (σ) for the noise component was determined based on the asynchrony observed between the two participants involved in the experiment. To quantify this asynchrony, we calculated the median of the absolute differences between the two participants for each trial, and then determined the interquartile range for each group and condition, resulting in a total of eight values of σ^62^.

In order to determine the optimal coupling weights for our study, we conducted an extensive analysis. Our implementation of the Kuramoto model was in accordance with previous study^13^. First, we systematically varied the values of *i*_1_, *i*_2_, *e*_1_, and *e*_2,_ ranging from 1 to 10 in increments of 1, resulting in 10,000 combinations. Within each combination, we simulated time-series data by employing different coupling matrices and initiated at random phases. Subsequently, we compared the resulting patterns of cross-correlation lag with those observed in the real data. Specifically, we focused on the patterns at lag-1, lag0, and lag+1 patterns, as these patterns provided valuable information and had been demonstrated to be effective in previous studies for parameter search^13^. To minimize the impact of random initial phases and noises, we repeat the simulations 200 times within each coupling combination and 250 sets of simulations for all the combinations. To obtain more precise coupling weights, we varied the values of *e*_1_, *i*_1_, *e*_2_, and *i*_2_ based on the best coupling weights simulated in step 1. We systematically adjusted these values by increments of -0.9 to +0.9, with a step size of 0.2. Additionally, we repeated this process 200 times for each combination and performed an overall parameter search 35 times. The combination of 250 iterations in step 1 and 35 iterations in step 2 was sufficient to achieve stable coupling weights. Consistent with previous study^13^, we computed the group-level (all behavioural data from all dyads were submitted into the modeling) instead of individual-level parameters to obtain stable results.

#### 4.5.3 fNIRS data preprocessing and analyses

*Preprocessing.* To ensure the integrity of our fNIRS time-series data, we first verified the temporal alignment of the two datasets, confirming that they had synchronized start and end times. Subsequently, we employed the Homer 2 toolbox to implement a range of preprocessing procedures^78^. We converted the raw optical intensity data to optical density using the *hmrIntensity2OD* function, followed by identification and removal of bad channels with the *enPruneChannels* function. We performed motion artifact correction, which involved identification and correction of motion artifacts using the *hmrMotionArtifactByChannel* and *hmrMotionCorrectSpline* functions. Next, we transformed the optical density data to concentration changes in HbO and HbR using the *hmrOD2Conc* function. Finally, a wavelet-based denoising method was applied to remove superficial physiological noise and related spurious connectivity from the data^79^.

*ROI construction.* Since fNIRS data are relative values, we standardized the HbO concentrations by converting them into *z*-scores using the mean and standard deviations of resting-state fNIRS data. We then proceeded to create 12 regions of interest (ROIs) based on the localizations of the fNIRS channels (**Fig. 1C**). Specifically, we averaged the channels that shared a common location to form each ROI. For precise channel locations and AAL labels, please refer to **Supplementary Table S3**. These ROIs were centered around various brain regions, namely: (1) right Auditory region (RA), (2) right Temporal-Parietal Junction (RTPJ), (3) right Motor region (RM), (4) right Dorsal-Lateral Prefrontal Cortex (RDLPFC), (5) right Superior Frontal Cortex (RSFC), (6) right Frontopolar (RF), (7) left Frontopolar (LF), (8) left Superior Frontal Cortex (LSFC), (9) left Dorsal-Lateral Prefrontal Cortex (LDLPFC), (10) left Motor region (LM), (11) left Temporal-Parietal Junction (LTPJ), (12) left Auditory region (LA).

*Brain activation. Z*-scores of the HbO concentrations were time-averaged within each ROI for each condition. This allowed us to obtain the brain activation values for each participant.

*Intra- and inter-brain synchronization.* Intra- and inter-brain synchronization was computed based on the wavelet transform coherence^80^. The wavelet transform coherence serves as a measure of correlation between signals x and y in the time-frequency plane, and is useful for analyzing nonstationary signals^81,82^. To calculate coherence values between each combination of ROIs, we used the *wcoherence* function in MATLAB 2022a. This allowed us to compute coherence values for different conditions based on the z-scores of HbO concentrations. The average coherence value between 0.01 and 0.04 Hz was calculated. This frequency band allowed us to exclude noise associated with cardiac pulsation (above 0.7 Hz), respiration (0.15 – 0.3Hz) and very low frequency fluctuations (below 0.01 Hz)^83^. Specifically, we computed coherence values for all combinations of the 12 ROIs with each other, resulting in a total of 12 ROIs * 12 ROIs for Participant A, Participant B, and the interaction between Participant A and B under the four conditions.

*Granger causality analysis (GCA).* To examine the directionality of brain synchronization, we employed GCA using the *MVGC* toolbox in MATLAB^84,85^. We computed the conditional GCA values of both directions (from Participant A to B and from Participant B to A) using state space models, which are shown to offer enhanced statistical power and reduced bias compared to pure autoregressive estimators^84^. Our GCA was based on preprocessed, z-transformed HbO signals during the task periods.

#### 4.5.4 Statistical analyses

The data underwent analyses using linear mixed-effects modeling (LMM), implemented with the *lme4* package in R^86^. To assess the significance of model parameters, we used the *lmerTest* package^87^. The model incorporated Group (2 levels: SCZ-HC group vs. HC-HC group) and Condition (4 levels: Hearing Self vs. Hearing Each Other vs. Hearing A vs. Hearing B), along with their interaction, as fixed effects. Random effects were estimated for Dyad ID.

Type III ANOVA with Satterthwaite’s method for degrees of freedom was performed on these models using the *anova* function from the *car* package^88^. Next, significance tests were conducted using Satterthwaite *p* values^89^, and partial eta squared (η_p_^2^) was used to computed effect size using the *effectsize* package^90^. In case where significant main effects or interaction were observed, follow-up contrasts were further performed using the *emmeans* package in R^91^, with a Tukey adjustment applied to control for multiple comparisons. For fNIRS data analyses, LMM was performed for each ROI.

If the data distribution departs from the assumption of normality, as determined by the Shapiro-Wilk test, we reported the median and median absolute deviation (MAD) as descriptive statistics instead of the mean and standard deviation; also, we used Pearson correlation to examine the relationship between the measures of multi-brain network (including IBS and intra-brain synchronization) and behavioural performance. A Rank-Based Inverse Normal Transformation (RIN) would be employed prior to using Pearson correlation due to non-normal data. RIN is a non-linear transformation method that approximates normality regardless of the initial distribution shape^92^. Nonlinear transformations could improve marginal normality and linearity while minimizing the impact of outliers^92^. For multivariate correlation, we used the mediation model to further explore their possible mechanisms. We used *PROCESS* v3.5 to build mediation model^93^.

Additionally, we employed Bayesian network analysis to examine directional dependencies among variables^28^. We used Bayesian networks to generate a directed acyclic graph structure, using the hill-climbing algorithm with reference to previous study on SCZ population^28^. Stability analyses were conducted to assess arc strength, representing confidence in edge frequency and direction. The Bayesian network structure was determined based on edge frequency in 1000 bootstrap networks, retaining edges appearing in at least 85% of networks and pointing in the direction in at least 51% of the networks^94^. The significance of edges was evaluated using the Bayesian Information Criterion (BIC), with lower values indicating greater importance^28^. We employed the *bnlearn* package for Bayesian network analysis^95^.

To identify the connectivity between two ROIs that closely matched the coupling weights, we employed the Mahalanobis distance method. The Mahalanobis distance quantifies the distance between a point (e.g., a group-level computational parameter) and a distribution (e.g., a distribution of individual-level IBS), considering the covariance structure of the data^96^. It measures the number of standard deviations a point deviates from the mean of the distribution, providing a scale-invariant and unitless distance metric that accounts for dataset correlations^97^. To calculate the Mahalanobis distances, we utilized the MATLAB function *mahal* from the Statistics Toolbox. Since only IBS significantly related to behaviour synchronization, our analysis focused solely on IBS associated with the external coupling weights.

To predict symptomatology based on IBS, we used Partial Least Squares Regression (PLSR)^98^. PLSR consists of two steps: dimensionality reduction and regression^98^. In the first step, samples are projected into a lower-dimensional subspace. In the second step, the targets (y) are predicted using the transformed X. PLSR is a supervised method, meaning it focuses on retaining the features that are most relevant for predicting the target, rather than the features with the highest variance^99^. The regression part of PLSR can be seen as a form of regularized linear regression, where the number of components determines the strength of the regularization^99^.

PLSR is well-suited for handling multi-variables and addressing multicollinearity among predictors, making it a suitable choice in similar studies conducted previously^100,101^. Similarly, we aimed to predict symptomatology dimensions, such as the total PANSS score and its four factors (negative, positive, affective, and cognitive symptoms), from a multi-brain network. Given that PLSR identifies the most relevant components from the original features, our interest lies in finding key contributors to the latent variables. To achieve this, we utilized the scikit-learn package to construct the PLSR model^99^. In instances where data were missing, pair-wise deletion was applied to the dataset, meaning that incomplete cases were excluded from the analyses.

## Data and Code Accessibility

All behavioural data and custom code for neural analysis have been made publicly available via the Open Science Framework and can be accessed at https://osf.io/gmjz7/. Further inquiries about research can be made to the corresponding authors.

## Author contributions

Y.P., J.C., and Z.L. conceived of the project and designed the experiments; Y.J.W., Y.W., and L. Z. implemented the experiments and collected data; Y.J.W. analysed the data under the supervision of Y.P. and J.C.; Y.P, J.C., Z.L., L.Z. and Y.J.W. interpreted the results; Y.J.W. wrote the original manuscript. Y.P, J.C. provided critical revisions. All authors approved the final version of the manuscript for submission.

## Acknowledgements

The authors would like to extend their sincere gratitude to Ivana Konvalinka and Ole Adrian Heggli for their invaluable guidance in Kuramoto modelling. Furthermore, the authors would like to thank Jie Pu, Chang Su and other assistants for their support in data collection. They also appreciate the support of all the staff in the Department of Psychiatry of the Second Affiliated Hospital, Zhejiang University School of Medicine, for their cooperation in conducting the psychological assessments.

