## Supplementary material for "A computational and multi-brain signature for aberrant social coordination in schizophrenia": Supplemtary Materials

**This PDF file includes:**

Supplementary Text

Figure S1

Figure S2

Figure S3

Figure S4

Table S1

Table S2

Table S3

Table S4

Table S5

Table S6

Table S7

Table S8

**Supplementary Text**

Subjective measurements

*Interpersonal Reactivity Index.* The Interpersonal Reactivity Index^1^ was developed as a comprehensive measurement tool to assess empathy across multiple dimensions instead of a singular construct. This self-report measure consists of 28 items, which are further divided into four subscales, each comprising seven items. These subscales include Perspective Taking, which assesses the tendency to adopt the psychological viewpoint of others; Fantasy, which captures the capacity to imaginatively immerse oneself in the emotions and actions of fictional characters; Empathic Concern, which measures the feelings of sympathy and compassion towards others; and Personal Distress, which evaluates the self-oriented experience of anxiety and unease in tense interpersonal settings.

*Schizophrenia Quality of Life Scale.* The Schizophrenia Quality of Life Scale has been established as a practical and widely accepted method for assessing self-reported quality of life in individuals with schizophrenia^2^. This comprehensive scale encompasses a total of 30 items, which are further categorized into three distinct subscales: Psychosocial, Motivation and Energy, and Symptoms and Side-effects. Within the psychosocial subscale, 15 items are dedicated to evaluating emotional expression and interpersonal communication. The motivation/energy subscale consists of 7 items that specifically gauge motivation levels and overall energy. The symptom/side effect subscale comprises 8 items, providing a comprehensive assessment of symptoms and potential drug-related side effects.

*Liebowitz Social Anxiety Scale.* The Liebowitz Social Anxiety Scale assesses the complete range of performance and social difficulties commonly reported by individuals with social phobia^3^. It effectively measures social anxiety levels by evaluating individuals' fear and avoidance during social interactions and performances. The scale consists of 24 items, with 13 items specifically addressing performance anxiety and 11 items focusing on social situations. Each item includes subscales for social fear and social avoidance, allowing for the assessment of varying levels of severity in social anxiety.

Apparatus

For the tapping task, we affixed two pressure sensor pads onto the table surface and connected them to a computer to record the timing of the taps. We designed four buttons that allowed experimenter to control the auditory feedback that participants received during the task. To ensure that participants could discern between the auditory stimuli, we selected distinctive frequencies for different auditory stimulus: 2093 Hz for the metronome, 131 Hz for role A, 234 Hz for role B, and 1000 Hz for the stop signal. The metronome and auditory feedback were designed to persist for a duration of 50 ms, providing a brief and consistent auditory cue for the participants. In contrast, the stop-signal lasted for 500 ms to ensure that participants clearly knew when to stop. To avoid registering two taps that occurred too closely together, we set a threshold of 150 ms between consecutive taps. Specifically, if two taps occurred within this time window, only the first tap was recorded.

Behavioral correlations in the HC-HC group

In the HC-HC group, we found that rhythm deviation was significantly related to tapping variability in Role B (*r* = 0.23, *p_FDR_* = 0.005) but not in Role A (*r* = 0.12, *p_FDR_* = 0.099). The rhythm deviation was negatively correlated with synchronization index in both roles: Role A: *r* = -0.22, *p_FDR_* = 0.007; Role B: *r* = -0.40, *p_FDR_* < 0.001 (**Supplementary Figure S1**).

Conditional differences

Eleven ROI combinations for inter-brain synchronization (IBS) exhibited significant condition differences (**Supplementary Table S5**). Estimated marginal means analysis further revealed that Hearing B demonstrated the highest IBS, which significantly differed from the other three conditions in all 11 ROIs (**Supplementary Figure S2**).

Group differences

For intra-brain synchronization, lower synchronization levels were observed in 9 ROI combinations in the SCZ-HC group and 4 ROI combinations in the SCZ individuals (**Supplementary Table S6** and **Figure S3**). Similarly, a reduction in intra-brain GCA was observed in 9 out of 11 combinations of ROIs in the SCZ-HC group (**Supplementary Table S7** and **Figure S4**). However, no significant conditional differences were observed in any intra-brain connectivity.

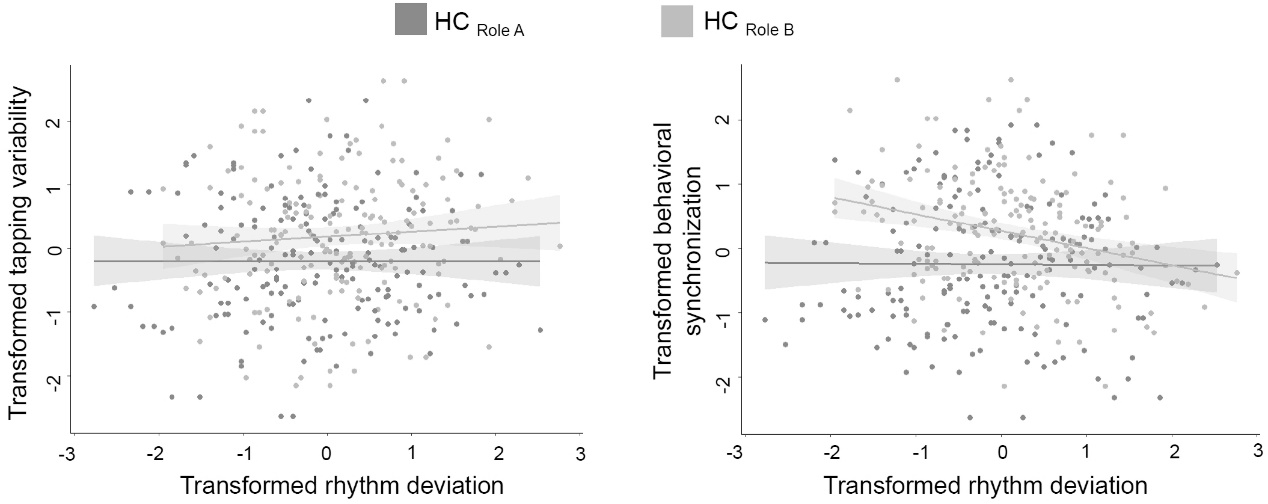

**Figure S1.** The scatter plots among behavioral indices within the HC-HC group. The data underwent a Rank-Based Inverse Normal Transform to conform a normal distribution.

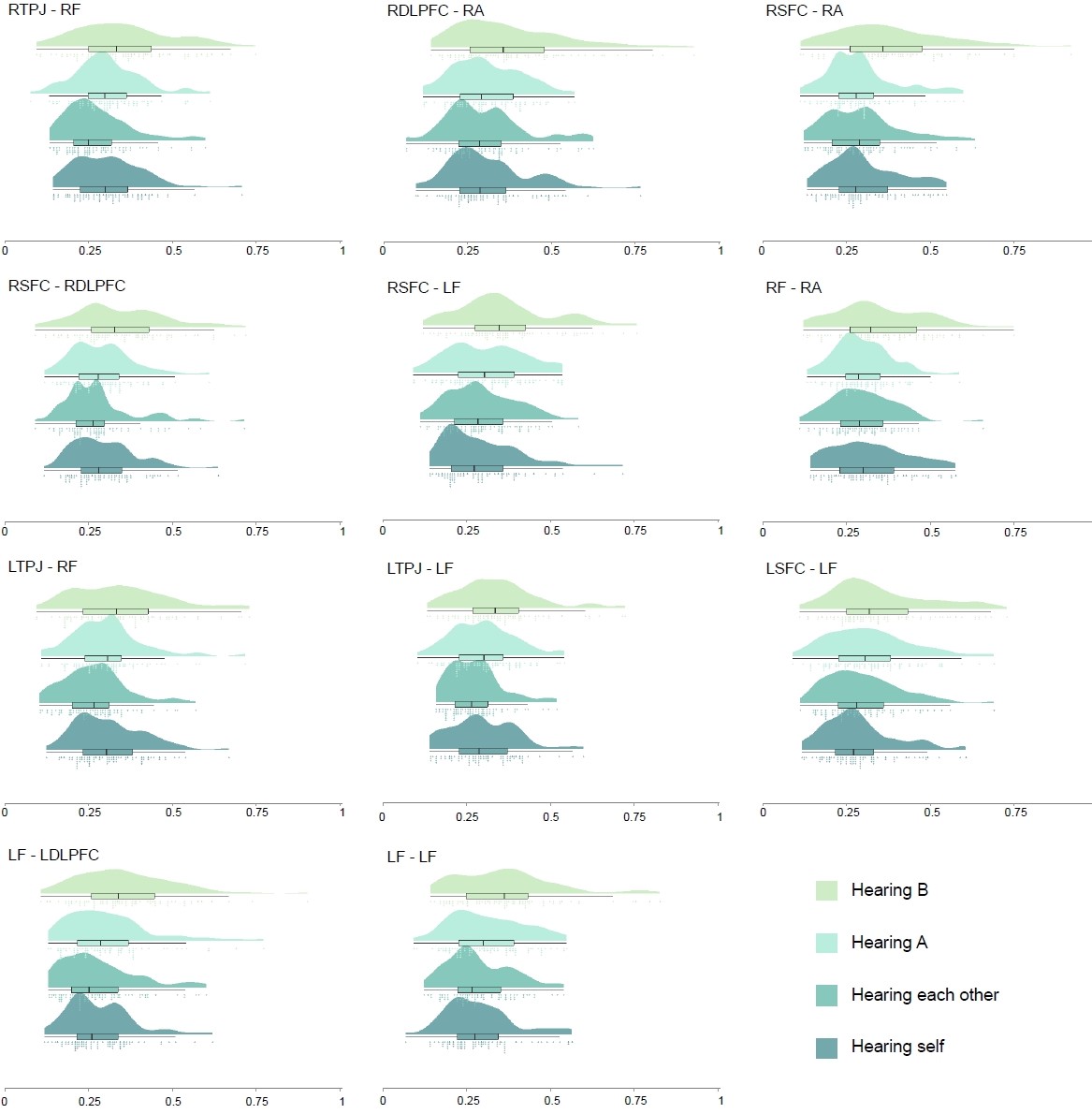

**Figure S2.** The data distribution for inter-brain synchronization (IBS) at different ROI combinations showing conditional differences. The x-axis indicates the IBS strength.

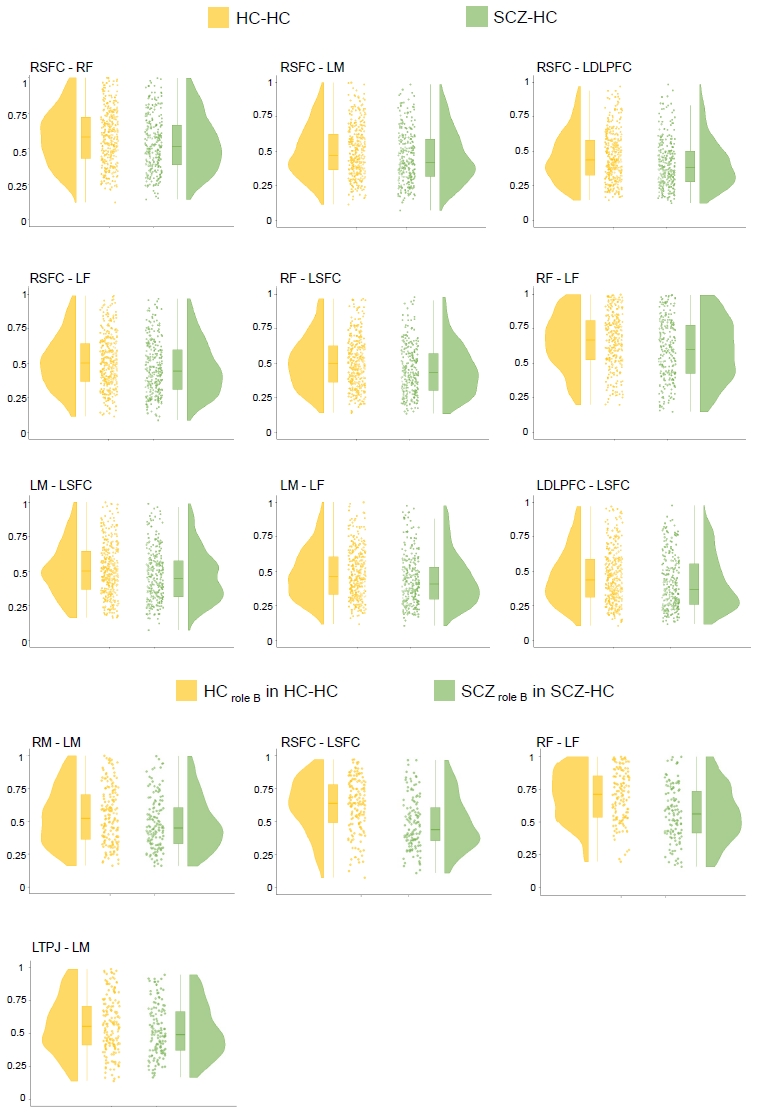

**Figure S3.** The data distribution for intra-brain synchronization at different ROI combinations showing group differences.

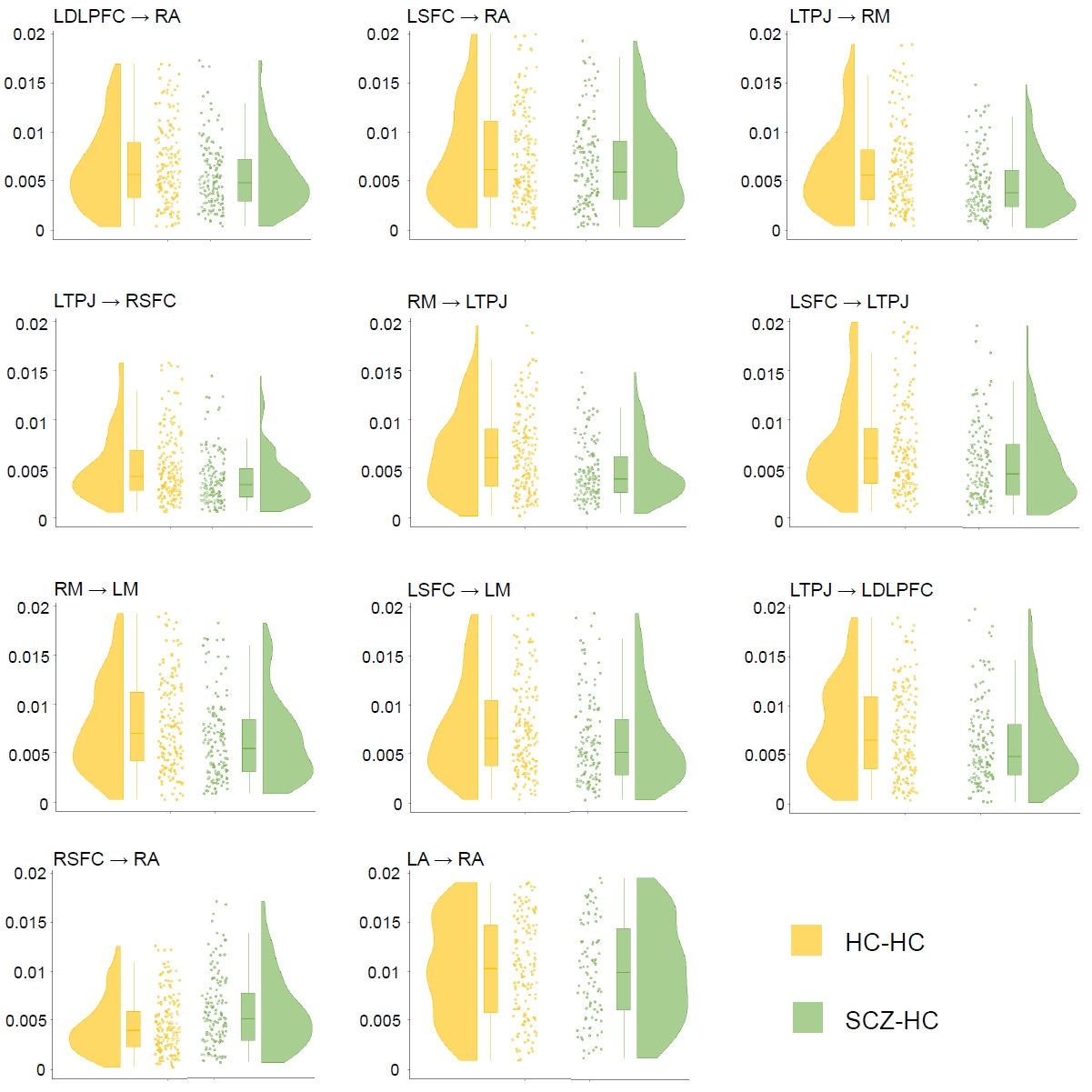

**Figure S4.** The data distribution for intra-brain Granger Causality at different ROI combinations showing group differences.

**Table S1. Demographic Data**

|  | SCZ | | HC | | *P* value (Mann-Whitney’ test) |
| --- | --- | --- | --- | --- | --- |
| Characteristic | Value | No. | Value | No. |  |
| **Demographic data** | | | | | |
| Age | 25.00 (7.41) | 35 | 23.00 (4.45) | 123 | 0.54 |
| Gender | Female | 19 | Female | 65 | 0.90 |
|  | Male | 16 | Male | 58 |  |
| **Clinical Interview** | | | | | |
| PANSS | | | | | |
| Total PANSS Score | 58 (16.3) | 33 | NA | NA | NA |
| Negative Symptom | 3.32 (1.14) | 33 | NA | NA | NA |
| Positive Symptom | 4.45 (2.16) | 33 | NA | NA | NA |
| Affective Symptom | 5.60 (2.52) | 33 | NA | NA | NA |
| Cognitive Symptom | 4.84 (1.91) | 33 | NA | NA | NA |
| FSQ-MIS | | | | | |
| Total score | NA | NA | 0.00 (0.00) | 110 | NA |
| **Self-report** | | | | | |
| Interpersonal Reactivity Index | | | | | |
| Fantasy Scale | 11.00 (4.45) | 30 | 14.00 (5.93) | 123 | 0.04 |
| Empathetic Concern | 15.50 (5.19) | 30 | 17.00 (2.97) | 123 | 0.03 |
| Perspective taking | 14.00 (5.19) | 30 | 18.00 (2.97) | 123 | <.001 |
| Personal distress | 15.50 (2.97) | 30 | 15.00 (2.97) | 123 | 0.83 |
| Schizophrenia Quality of Life Scale | | | | | |
| Psychosocial | 0.22 (0.25) | 30 | 0.15 (0.10) | 123 | 0.02 |
| Motivation and Energy | 0.50 (0.13) | 30 | 0.61 (0.11) | 123 | 0.001 |
| Symptoms and Side-effects | 0.13 (0.14) | 30 | 0.06 (0.09) | 123 | 0.004 |
| Liebowitz Social Anxiety Scale | | | | | |
| Social Fear | 12.50 (13.30) | 30 | 15.00 (10.40) | 123 | 0.48 |
| Social Avoidance | 15.00 (17.80) | 30 | 20.00 (11.90) | 123 | 0.45 |

Notes: Report the value of results with median (MAD) format; NA, not applicable.

**Table S2. Pairwise Demographic Data**

|  | SCZ-HC dyads | | HC-HC dyads | | | *P* value (Mann-Whitney’ test) |
| --- | --- | --- | --- | --- | --- | --- |
| Characteristic | Value | No. | | Value | No. |  |
| Pair age | 24.50 (7.41) | 35 | | 22.50 (3.71) | 44 | 0.24 |
| Within-pair age difference | 1.00 (1.48) | 35 | | 1.00 (1.48) | 44 | 0.25 |
| Pair gender | Female-Female | 13 | | Female-Female | 15 | 0.63 |
|  | Female-Male | 13 | | Female-Male | 15 |  |
|  | Male-Male | 9 | | Male-Male | 14 |  |

Notes: Report the value of results with median (MAD) format; NA, not applicable.

**Table S3**. Channel locations and AAL labels

| ROI | Channel | x | y | z | AAL Label |
| --- | --- | --- | --- | --- | --- |
| Right Auditory region | 1 | 70.95 | -32.78 | -15.54 | Temporal_Mid_R |
|  | 2 | 71.05 | -45.48 | -3.05 | Temporal_Mid_R |
|  | 4 | 67.99 | -6.49 | -1.41 | Temporal_Sup_R |
|  | 8 | 69.91 | -17.36 | 10.58 | Temporal_Sup_R |
| Right Temporal-parietal Junction | 3 | 66.88 | -56.98 | 5.48 | Temporal_Mid_R |
|  | 6 | 65.05 | -57.15 | 25.72 | Angular_R |
|  | 7 | 60.93 | -57.21 | 42.00 | Parietal_Inf_R |
| Right Motor region | 11 | 63.91 | -2.25 | 35.62 | Postcentral_R |
|  | 13 | 57.10 | -28.45 | 55.08 | Parietal_Inf_R |
|  | 14 | 56.89 | -1.78 | 50.77 | Precentral_R |
|  | 15 | 44.67 | -3.06 | 61.23 | Frontal_Mid_R |
| Right Dorsal-lateral Prefrontal Cortex | 12 | 55.84 | 23.30 | 34.51 | Frontal_Inf_Oper_R |
|  | 16 | 47.72 | 41.21 | 27.90 | Frontal_Mid_R |
| Right Superior Frontal Cortex | 19 | 24.51 | 51.12 | 40.87 | Frontal_Sup_R |
|  | 20 | 12.68 | 40.56 | 55.30 | Frontal_Sup_R |
| Right Frontopolar | 17 | 38.29 | 58.16 | 20.45 | Frontal_Mid_R |
|  | 21 | 26.13 | 69.85 | 8.41 | Frontal_Sup_R |
|  | 22 | 13.86 | 72.29 | -8.26 | Frontal_Med_Orb_R |
|  | 23 | 14.23 | 64.53 | 29.03 | Frontal_Sup_R |
| Left Frontopolar | 26 | -11.78 | 65.23 | 30.23 | Frontal_Sup_Medial_L |
|  | 30 | -10.07 | 72.30 | -7.12 | Frontal_Sup_Orb_L |
|  | 31 | -23.53 | 69.89 | 11.16 | Frontal_Sup_L |
|  | 35 | -36.00 | 57.94 | 23.35 | Frontal_Mid_L |
| Left Superior Frontal Cortex | 27 | -11.96 | 40.43 | 55.92 | Frontal_Sup_L |
|  | 28 | -21.95 | 51.20 | 42.24 | Frontal_Sup_L |
| Left Dorsal-lateral Prefrontal Cortex | 36 | -45.93 | 41.38 | 31.89 | Frontal_Mid_L |
|  | 45 | -54.25 | 21.65 | 39.01 | Frontal_Mid_L |
| Left Motor region | 32 | -40.21 | -3.29 | 64.04 | Precentral_L |
|  | 33 | -54.62 | -28.41 | 59.24 | Postcentral_L |
|  | 34 | -51.17 | -4.78 | 56.15 | Precentral_L |
|  | 40 | -60.78 | -5.52 | 43.88 | Postcentral_L |
| Left Temporal-parietal Junction | 37 | -60.33 | -54.49 | 45.30 | Parietal_Inf_L |
|  | 38 | -65.60 | -55.29 | 27.76 | SupraMarginal_L |
|  | 46 | -67.23 | -55.58 | 10.38 | Temporal_Mid_L |
| Left Auditory region | 42 | -69.21 | -18.62 | 19.22 | Postcentral_L |
|  | 44 | -67.87 | -9.12 | 6.29 | Temporal_Sup_L |
|  | 47 | -70.92 | -43.84 | 3.81 | Temporal_Mid_L |
|  | 48 | -72.10 | -31.38 | -6.77 | Temporal_Mid_L |
| Uncategorized channels | 5 | 64.56 | 9.15 | 24.28 | Precentral_R |
|  | 9 | 70.12 | -29.66 | 23.33 | SupraMarginal_R |
|  | 10 | 68.11 | -29.76 | 40.47 | SupraMarginal_R |
|  | 18 | 35.82 | 34.93 | 48.53 | Frontal_Mid_R |
|  | 24 | -0.04 | 52.67 | 43.61 | Frontal_Sup_Medial_L |
|  | 25 | 2.37 | 68.88 | 11.65 | Frontal_Sup_Medial_R |
|  | 29 | -32.39 | 33.19 | 50.88 | Frontal_Mid_L |
|  | 39 | -65.80 | -29.75 | 47.04 | SupraMarginal_L |
|  | 41 | -70.26 | -29.62 | 29.37 | SupraMarginal_L |
|  | 43 | -64.09 | 4.96 | 32.32 | Precentral_L |

Table S4. The four dimensions of Positive and Negative Syndrome Scale and the items they contained.

| Dimensions | Item |
| --- | --- |
| Negative | Blunted affect |
|  | Emotional withdrawal |
|  | Poor rapport |
|  | Apathetic social withdrawal |
|  | Low Spontaneity / flow |
|  | Mannerisms and posturing |
|  | Motor retardation |
| Positive | Delusions |
|  | Hallucinations |
|  | Grandiosity |
|  | Unusual thought content |
| Affective | Suspiciousness / Persecution |
|  | Somatic concern |
|  | Anxiety |
|  | Guilt feelings |
|  | Tension |
|  | Depression |
|  | Active social avoidance |
| Cognitive | Conceptual disorganization |
|  | Hyperactivity / Excitement |
|  | Hostility |
|  | Difficulty in abstract thinking |
|  | Stereotyped thinking |
|  | Uncooperativeness |
|  | Disorientation |
|  | Poor attention |
|  | Lack of judgment and insight |
|  | Disturbance of volition |
|  | Poor impulse control |
|  | Preoccupation |

**Table S5**. ROI combinations of inter-brain synchronization showing conditional differences.

|  | Hearing self | Hearing each other | Hearing A | Hearing B | Conditional difference |
| --- | --- | --- | --- | --- | --- |
| RTPJ-RF | 0.30 (0.11) | 0.25 (0.08) | 0.30 (0.09) | 0.33 (0.13) | *F* = 7.12, *p*_fdr_ = 0.008, η_p_^2^ = 0.07 |
| RDLPFC-RA | 0.29 (0.10) | 0.29 (0.10) | 0.29 (0.11) | 0.36 (0.16) | *F* = 6.52, *p*_fdr_ = 0.012, η_p_^2^ = 0.07 |
| RSFC-RA | 0.28 (0.10) | 0.29 (0.10) | 0.28 (0.08) | 0.36 (0.15) | *F* = 10.19, *p*_fdr_ < 0.001, η_p_^2^ = 0.10 |
| RSFC-RDLPFC | 0.28 (0.09) | 0.26 (0.07) | 0.28 (0.09) | 0.33 (0.13) | *F* = 6.43, *p*_fdr_ = 0.012, η_p_^2^ = 0.07 |
| RSFC-LF | 0.27 (0.11) | 0.28 (0.11) | 0.30 (0.12) | 0.35 (0.12) | *F* = 6.43, *p*_fdr_ = 0.009, η_p_^2^ = 0.07 |
| RF-RA | 0.30 (0.12) | 0.29 (0.10) | 0.29 (0.08) | 0.32 (0.13) | *F* = 5.42, *p*_fdr_ = 0.041, η_p_^2^ = 0.06 |
| LTPJ-RF | 0.30 (0.11) | 0.27 (0.09) | 0.31 (0.08) | 0.33 (0.15) | *F* = 7.09, *p*_fdr_ = 0.008, η_p_^2^ = 0.07 |
| LTPJ-LF | 0.29 (0.11) | 0.27 (0.07) | 0.30 (0.11) | 0.34 (0.10) | *F* = 7.86, *p*_fdr_ = 0.005, η_p_^2^ = 0.08 |
| LSFC-LF | 0.27 (0.09) | 0.28 (0.10) | 0.30 (0.11) | 0.32 (0.12) | *F* = 6.46, *p*_fdr_ = 0.012, η_p_^2^ = 0.07 |
| LF-LDLPFC | 0.26 (0.09) | 0.25 (0.10) | 0.29 (0.12) | 0.34 (0.15) | *F* = 9.07, *p*_fdr_ = 0.001, η_p_^2^ = 0.09 |
| LF-LF | 0.27 (0.10) | 0.27 (0.11) | 0.30 (0.12) | 0.36 (0.13) | *F* = 9.94, *p*_fdr_ < 0.001, η_p_^2^ = 0.10 |

**Table S6**. ROI combinations of intra-brain synchronization showing group differences and group*role interaction effects.

|  | HC-HC | SCZ-HC | Group difference |
| --- | --- | --- | --- |
| RSFC-RF | 0.59 (0.23) | 0.52 (0.21) | *F* = 10.44, *p*_fdr_ = 0.029, η_p_^2^ = 0.02 |
| RSFC-LM | 0.48 (0.20) | 0.42 (0.18) | *F* = 10.46, *p*_fdr_ = 0.029, η_p_^2^ = 0.02 |
| RSFC-LDLPFC | 0.44 (0.19) | 0.38 (0.17) | *F* = 16.51, *p*_fdr_ = 0.007, η_p_^2^ = 0.03 |
| RSFC-LF | 0.51 (0.21) | 0.45 (0.21) | *F* = 12.71, *p*_fdr_ = 0.015, η_p_^2^ = 0.02 |
| RF-LSFC | 0.51 (0.21) | 0.43 (0.20) | *F* = 15.15, *p*_fdr_ = 0.007, η_p_^2^ = 0.02 |
| RF-LF | 0.74 (0.30) | 0.66 (0.33) | *F* = 11.48, *p*_fdr_ = 0.025, η_p_^2^ = 0.02 |
| LM-LSFC | 0.50 (0.21) | 0.45 (0.20) | *F* = 13.47, *p*_fdr_ = 0.013, η_p_^2^ = 0.02 |
| LM-LF | 0.47 (0.20) | 0.41 (0.18) | *F* = 10.43, *p*_fdr_ = 0.029, η_p_^2^ = 0.02 |
| LDLPFC-LSFC | 0.45 (0.21) | 0.37 (0.20) | *F* = 15.96, *p*_fdr_ = 0.007, η_p_^2^ = 0.03 |
|  | HC in  HC-HC | SCZ in  SCZ-HC | Group × Role interaction |
| RM-LM | 0.56 (0.28) | 0.45 (0.20) | *F* = 11.16, *p*_fdr_ = 0.046, η_p_^2^ = 0.02 |
| RSFC-LSFC | 0.66 (0.23) | 0.48 (0.22) | *F* = 16.08, *p*_fdr_ = 0.01, η_p_^2^ = 0.03 |
| RF-LF | 0.79 (0.28) | 0.60 (0.28) | *F* = 12.90, *p*_fdr_ = 0.02, η_p_^2^ = 0.03 |
| LTPJ-LM | 0.57 (0.24) | 0.49 (0.22) | *F* = 10.87, *p*_fdr_ = 0.046, η_p_^2^ = 0.02 |

**Table S7**. ROI combinations of intra-brain Granger Causality showing group differences.

| From | To | HC-HC | SCZ-HC | Group difference |
| --- | --- | --- | --- | --- |
| LDLPFC | RA | 0.0059 (0.0045) | 0.0048 (0.0031) | *F =* 16.03*, p_fdr_ =* 0.017*,* η*_p_^2^ =* 0.05 |
| LSFC | RA | 0.0070 (0.0062) | 0.0059 (0.0044) | *F =* 20.76, *p_fdr_ =* 0.0031*,* η*_p_^2^ =* 0.07 |
| LTPJ | RM | 0.0057 (0.0039) | 0.0038 (0.0027) | *F =* 23.15*, p_fdr_ =* 0.0020*,* η*_p_^2^ =* 0.08 |
| LTPJ | RSFC | 0.0042 (0.0027) | 0.0033 (0.0022) | *F = 15.97, p_fdr_ =* 0.017*,* η*_p_^2^ =* 0.05 |
| RM | LTPJ | 0.0061 (0.0043) | 0.0039 (0.0023) | *F = 22.03, p_fdr_ =* 0.0023*,* η*_p_^2^ =* 0.07 |
| LSFC | LTPJ | 0.0062 (0.0042) | 0.0045 (0.0036) | *F = 18.95, p_fdr_ =* 0.0052*,* η*_p_^2^ =* 0.06 |
| RM | LM | 0.0074 (0.0055) | 0.0054 (0.0038) | *F = 19.82, p_fdr_ =* 0.0041*,* η*_p_^2^ =* 0.07 |
| LSFC | LM | 0.0072 (0.0053) | 0.0052 (0.0042) | *F = 14.04, p_fdr_ =* 0.030*,* η*_p_^2^ =* 0.05 |
| LSFC | LDLPFC | 0.0074 (0.0061) | 0.0048 (0.0033) | *F = 26.88, p_fdr_ <* 0.001*,* η*_p_^2^ =* 0.09 |
| RSFC | RA | 0.0039 (0.0027) | 0.0050 (0.0033) | *F* = 14.83, *p*_fdr_ = 0.024, η_p_^2^ = 0.05 |
| LA | RA | 0.013 (0.0085) | 0.015 (0.011) | *F* = 14.62, *p*_fdr_ = 0.025, η_p_^2^ = 0.05 |

**Table S8.** ROI combinations of inter-brain synchronization that best represent external couplings derived from dynamical systems modeling.

| Group | Condition | *e*_1_ | *e*_2_ |
| --- | --- | --- | --- |
| HC-HC | Hearing Self | RTPJ-RSFC | RTPJ-RSFC |
|  | Hearing Each Other | RM-LM | RM-LM |
|  | Hearing HC _Role A_ | RDLPFC-RTPJ | RDLPFC-RTPJ |
|  | Hearing HC _Role B_ | RSFC-RA | RSFC-RA |
| SCZ-HC | Hearing Self | LF-LF | RDLPFC-LTPJ |
|  | Hearing Each Other | LSFC-RSFC | LSFC-RSFC |
|  | Hearing HC _Role A_ | LA-RTPJ | RDLPFC-LSFC |
|  | Hearing SCZ _Role B_ | RSFC-RA | RSFC-RA |
